# A molecular circuit regulates fate plasticity in emerging and adult AT2 cells

**DOI:** 10.1101/2025.04.28.650846

**Authors:** Amitoj S Sawhney, Brian J Deskin, Junming Cai, Daniel Gibbard, Gibran Ali, Annika Utoft, Xianmei Qi, Aaron Olson, Hannah Hausman, Liberty Sabol, Shannon Holmberg, Ria Shah, Rachel Warren, Stijn De Langhe, Rui Benedito, Zintis Inde, Kristopher A Sarosiek, Evan Lemire, Adam L Haber, Liu Wang, Zong Wei, Douglas G Brownfield

## Abstract

Alveolar AT1 and AT2 cells are vital for lung gas exchange and become compromised in several diseases. While key differentiation signals are known, their emergence and fate plasticity are unclear. Here we show in the embryonic lung that single AT2s emerge at intermediate zones, extrude, and connect with nearby epithelium via interlumenal junctioning. We observe that AT2s retain fate plasticity until the bZIP transcription factor C/EBPα suppresses Notch signaling at a novel *Dlk1* enhancer. Both *Dlk1* and *Cebpa* are regulated by the polycomb repressive complex (PRC2), which together form a “pulse generator” circuit that times *Dlk1* expression and thus Notch activation, resulting in a “salt and pepper” pattern of AT1 and AT2 fate. In injured adult lungs, C/EBPα downregulation is required to re-access AT2 fate plasticity and is mediated by the dominant negative C/EBP family member CHOP. Finally, *Cebpa* loss also activates a “defender” AT2 state, distinct from its reparative state, and we propose AT2s toggle between either state following infection to protect and repair alveoli.

## Introduction

Alveoli are the sites of gas exchange in the lung that are lined by two distinct epithelial cell types. The flat surface of alveolar type 1 (AT1) cells facilitates gas exchange while cuboidal alveolar type 2 (AT2) cells secrete surfactant to prevent alveolar collapse. Defective alveoli underly several maladies—bronchopulmonary dysplasia (BPD) in infants, chronic obstructive pulmonary disease (COPD)/emphysema^1^, pulmonary fibrosis^2^, lung adenocarcinoma^3, 4^, and respiratory infections like Severe Acute Respiratory Syndrome (SARS)^5, 6^ and COVID-19^7^.

Deciphering how these crucial alveolar cell types emerge as well as the mechanisms that regulate their fate is imperative for developing therapies that will not only slow or stop disease progression but may even promote lung repair.

As alveologenesis begins, AT1 and AT2 cells arise from a common pool of distal progenitors (DP)^4, 8, 9, 10, 11^, with differentiation detected as early as embryonic day 16.5 (e16.5) in mice, save for one study reporting an earlier timepoint^12^. Upon differentiation, AT1 and AT2 cells are arranged in a salt and pepper pattern^13^ along a lumenal surface which, as alveoli mature, interconnect with lumens of anatomically distinct branches to facilitate gas conduction between alveoli^14^. After development, mature AT2 cells can act as antigen presenting cells in an immune response^15, 16^ (partly directed interferon signaling^17, 18^), as well as facultative stem cells following injury^4, 11, 19, 20^ ^21^ (directed by Wnt^22^ and Notch^23^ signaling). While both AT2 states are critical after infection for alveolar recovery, a recent study found interferon stimulation suppresses repair of the lung epithelium following infection^24, 25^, suggesting that signals driving these states may oppose one another at the cellular level.

Here we show nascent AT2 cells are first detected molecularly at e15.5 as singletons at stereotyped intermediate regions of distal branches. Following basal extrusion, we observe nascent AT2s can form *de novo* attachments to nearby but anatomically separate epithelia, a step that is likely critical in stitching together alveolar lumen and that we term “interlumenal junctioning”, followed by sporadically patterned AT2 maturation, distinct from the “proximal-to-distal wave” pattern known to initiate differentiation of AT1s and AT2s earlier^13^.

We determine that nascent AT2s retain fate plasticity into the first perinatal week, revising the lineage model to include a “window of fate plasticity” spanning from distal progenitors to the AT2 lineage. From screening transcriptional regulators around this window, we identify a critical negative regulator of AT2 fate plasticity, the bZIP transcription factor C/EBPα, that when lost results in AT2-to-AT1 conversion. Previously C/EBPα has been reported to play a role in epithelial homeostasis^26^ and is implicated in fate regulation in multiple tissues, including the blood and liver^27^. Further, we found that the observed AT2-to-AT1 differentiation was mediated in a non-cell autonomous manner by the Notch regulator DLK1, a cell-surface transmembrane protein^28^ that has been associated with brain and muscle development, stem cell maintenance, lung cancer, and abnormal tissue repair^23, 29^. Observing a pulsed expression of *Dlk1* during alveologenesis, we find *Dlk1* and *Cebpa* are both regulated by the polycomb repressive complex (PRC2), and determine these components together constitute an incoherent feed forward loop. Upon abrogation of this “pulsed generator” circuit, we observe a disruption of AT1/AT2 differentiation. PRC2 is a well-established transcriptional repressor that has been shown to regulate DLK1 expression in neuronal fate selection^30^ and recent evidence suggests it also modulates C/EBPα expression in the context of adipocyte differentiation^31^.

Following injury in the adult lung C/EBPα must be downregulated in AT2s for conversion to AT1. We identify that the dominant negative member of the C/EBP family, CHOP, is induced in AT2s following injury and its upregulation promotes AT2-to-AT1 differentiation. Finally, we observe *Cebpa* loss also activates a “defensive” AT2 state that is enriched in antipathogen response genes, likely regulated by interferon signaling, and is mutually exclusive from the AT1-associated DATP state^32^.

## Results

### Timing and pattern of AT2 cell emergence

To delineate newly forming AT2 cells from distal progenitors and their mature counterparts, we analyzed scRNAseq data of the distal epithelial lineage from a prior study sampling the embryonic, perinatal, and adult timepoints of mouse lung development^33^ (Fig. 1a). Clustering analysis of the distal epithelium identified both DP and AT1 populations, as well as stratified AT2s into two groups (Fig. 1b) that are also largely separate in time (Fig. 1c). While these AT2 groups positively correlate in gene expression (Pearson coefficient = 0.6, Suppl. Fig. 1a), their distinction as a separate cluster was confirmed by co-assignment probability (Suppl. Fig. 1b).

**Figure 1:**
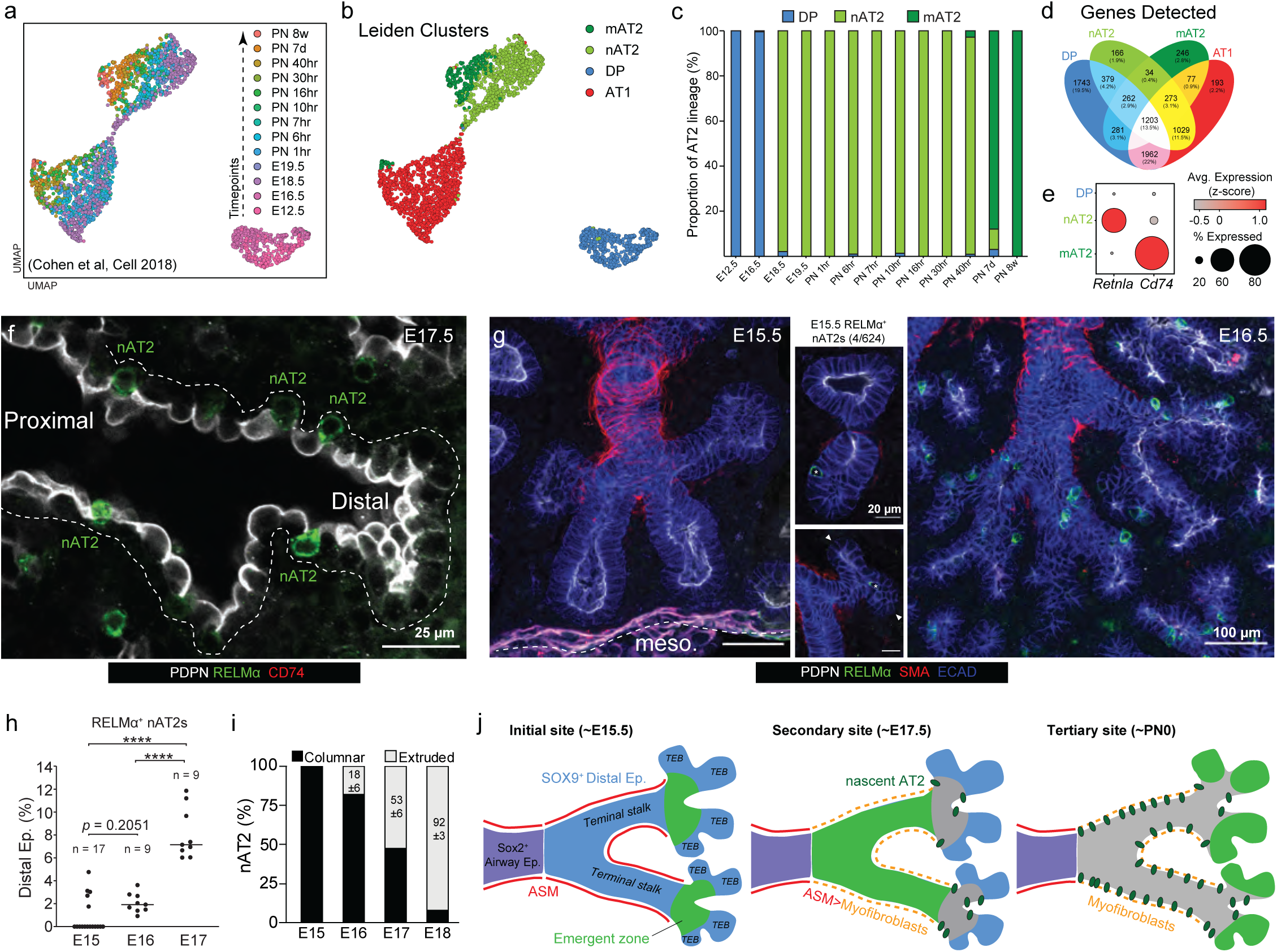
Emergence and patterning of nascent AT2 cells. **a** scRNAseq data (GSE119228)^33^ depicting embryonic, perinatal, and adult timepoints of alveolar epithelial development. **b** Leiden Clustering analysis identifies four populations: mature AT2s (mAT2), nascent AT2s (nAT2), distal progenitors (DP), and AT1s. **c** Stratification of the AT2 lineage clusters across development depicting time separation of the DP, nAT2, and mAT2 cell populations. **d** Venn diagram showing number of genes detected for all clusters. **e** Dot plot showing stage-restricted expression of selected surface markers for nAT2 and mAT2 cells, *Retnla* and *Cd74* respectively. **f** e17.5 lung immunostained for nascent AT2 marker RELMα (green), mature AT2 marker CD74 (red), and DP/AT1-restrictive luminal marker PDPN (white). Bar, 25 µm. **g** e15.5 (left) and e16.5 (right) lungs immunostained for ASM cell marker smooth muscle actin (SMA, red), RELMα (green), epithelial marker E-cadherin (ECAD, blue), and PDPN (white). Close-up images for e15.5 lungs depict rare cases of single nAT2 cells (asterisk) emerging in intermediate regions, but not at the terminal end buds (arrowheads). Bars, 100 µm (left and right panels), 20 µm (close-ups). Mesothelium marked with white dash. **h** Quantification of RELMα^+^ nAT2 emergence in distal epithelial branches across development from e15.5 to e17.5 determined by immunostaining (*n* represents branches scored for each timepoint). \*\*\*\**p* ≤ 0.0001 (Brown-Forsythe and Welch ANOVA test). **i** Quantification of RELMα^+^ nAT2 anatomical positioning (columnar versus basally extruded) across development. Extruded values presented as mean ± SD (*n* represents cells scored at each timepoint). **j** Schematic showing region of AT2 Emergent Zone (green) within Sox9^+^ distal epithelium at the terminal stalk (blue) instead of Sox2^+^ airway epithelium region. Around 17.5, nAT2 cell (RELMα^+^) basally extrude and then junction in distal epithelial branches and propagate emergent zone proximally further. Near birth (PN0), as proximal regions further developed, emergent zones shift to TEB.

Taken together with the temporal pattern, we refer to the earlier cluster as the nascent AT2 (nAT2) state, and the later as the mature AT2 (mAT2) state. From this distinction, we were able to identify hundreds of genes unique to either DP, nAT2, or mAT2 cluster (Fig. 1d, Suppl. Fig. 1c, and Suppl. Table 2), and initially focused on two for protein-level validation: *Retnla* expressed by nAT2s and *Cd74* by mAT2s (Fig. 1e, Suppl. Fig. 1d). *Retnla* encodes a regulator of inflammation, resistin-like molecule (RELM) α, while CD74 is critical in MHC-II antigen processing. In contrast to commonly used markers that label both DP and AT2 lineage starting around e17.5 (such as SFTPC)^4, 10^, RELMα uniquely labels nAT2s at this stage and can be used to distinguish them from both DPs and AT1s, as can be seen at an e17.5 transition zone where the mAT2 marker CD74 is not highly expressed (Fig. 1f, Suppl. Fig. 1e). Further, flow cytometry confirmed at e18.5 that a substantial proportion of marker-expressing AT2s are RELMα⁺ CD74⁻ (Suppl. Fig. 1g), which shifts to RELMα^-^ CD74^+^ in adult AT2s (Suppl. Fig. 1h).

Given the distinct labeling of nAT2s by RELMα in the embryonic lung, we used it to identify when and where AT2 cells first emerge during alveologenesis. Careful immunostaining revealed nAT2s emerge as early as e15.5 (Fig. 1g), which steadily increases over time (Fig. 1h). At first emergence, SOX9^low^/RELMα^+^ and nAT2s are singletons (Suppl. Fig. 1e, f) with retained columnar morphology (Fig. 1i) positioned at anatomical intermediate zones, located between airway smooth muscle (ASM)-covered distal branch stalks and terminal end buds (Fig. 1g, asterisks). After their initial emergence, nAT2s next arise (∼e17.5) within distal stalks following ASM to myofibroblast remodeling as previously reported^34^, then finally at terminal end buds soon after birth^4^. These observations support a sequential, three zone model of AT2 differentiation (Fig. 1j, left, Suppl. Fig. 1d, e) that integrates previously conflicting findings that reported AT2 differentiation first occurring either at terminal end buds^12^ or in a proximal-to-distal pattern^4, 9^.

### Nascent AT2s junction with nearby but anatomically distinct epithelium

While imaging RELMα at these timepoints, we also observed nAT2s forming connections to lumens across anatomically separate branches (Fig. 2a), a behavior which increased in frequency over time (Fig. 2b). While it is known that during alveolar maturation AT2 cells are capable of simultaneously integrating into lumens of multiple alveoli^14, 34^, the process by which it occurs has yet to be described. Careful immunostaining of junctioning nAT2s revealed their negativity for AT1 markers (PDPN, RAGE, and HOPX) and positivity for AT2 markers (SFTPC, LAMP1, and MUC1) that is often higher than non-junctioned AT2s, suggesting they may be further along in differentiation (Suppl. Fig. 2a, b). Using high resolution imaging and 3D reconstruction, we observed that junctions occurred either directly with the second lumen, or indirectly by attachment to another extruded nAT2 (Fig. 2c, Suppl. Fig. 2c, Suppl. Video 1-4).

**Figure 2:**
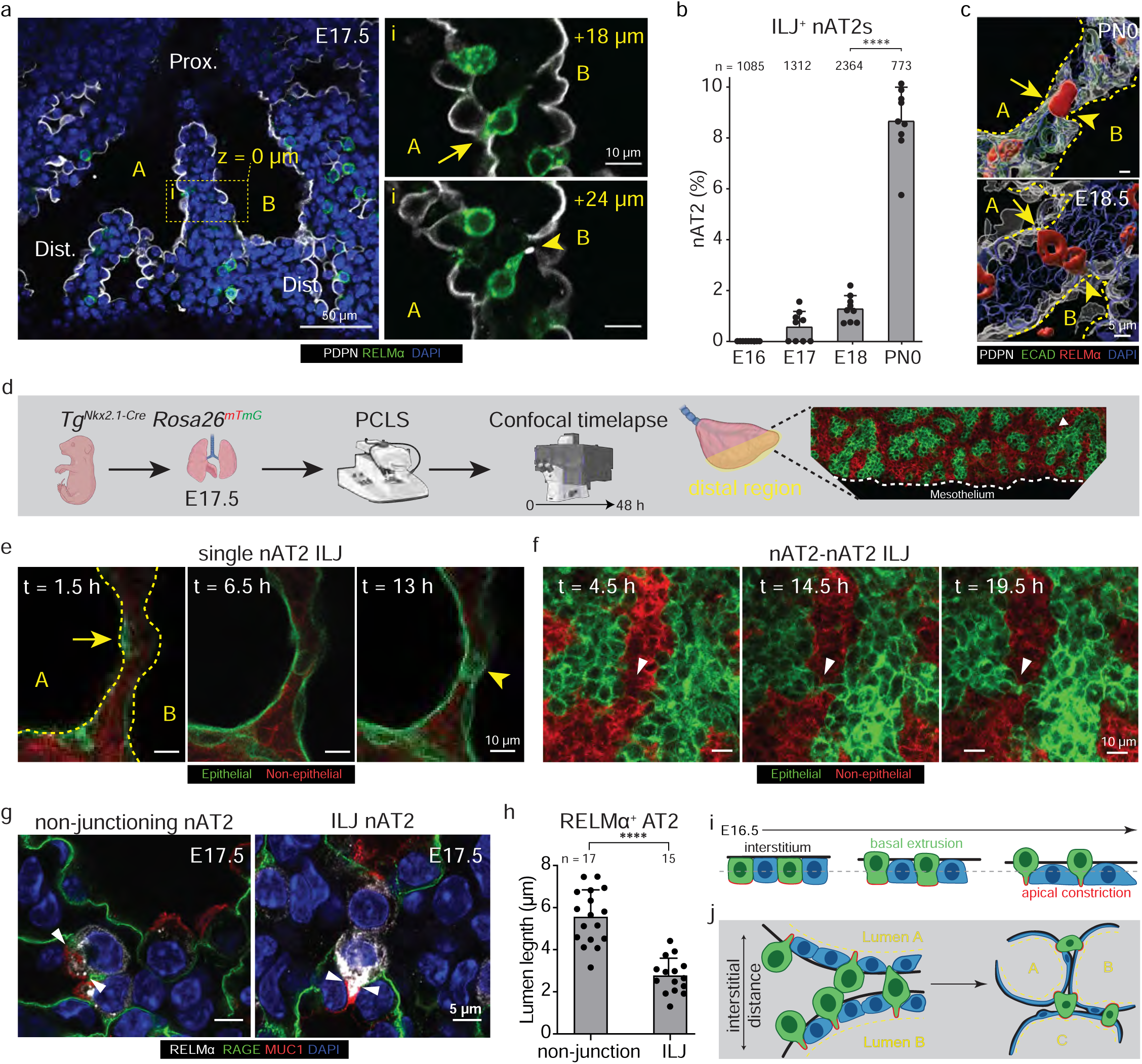
**a** e17.5 lung immunostained for RELMα (green), PDPN (white), and DAPI (blue). **a-i** depicts a RELMα^+^ nAT2 cell traversing across the interstitium from lumen A to B. Close-up panels (right) show a nAT2 cell embedded in lumen A, at z-position +18 µm (top right), as well as in lumen B, at z-position +24 µm (bottom right). Bar, 50 µm (left), 10 µm (close-ups; right). **b** Quantification of nAT2 interlumenal junctioning (ILJ) events detected during development by immunostaining (*n* represents nAT2 cells scored at each timepoint). \*\*\*\**p* = 4.8 × 10^-11^ (one-way ANOVA, data as mean ± SD). **c** 3D rendering immunostaining of single (upper) and double (lower) interlumenal junctioning nAT2s for E-cadherin (green), RELMα (red), PDPN (white), and DAPI. Lumen boundaries are marked in yellow dashes. **d** experimental schematics of timeplase confocal microscopy on *Nkx2.1^Cre^; Rosa26^mTmG^* lineage labeled precision cut lung slices at distal lung. **e** Single nAT2 junctions between two lumens in 12-hour snapshots. **f** Two nAT2s established contact at 14.5 hr and junction between two lumens in 15-hour snapshots, contact marked with white arrows. **g** Examples of luminal Muc1 domain length in non-junctioning (left) and junctioning nAT2s **h** Quantification of (g) shows condensed MUC1 domain in ILJ nAT2s. (*n* represents number of nAT2 cells scored at each timepoint). \*\*\*\**p* = 4.8 × 10^-11^ (Student’s two-sided *t*-test, data as mean ± SD). **i** schematics of stages of interluminal junctioning, including nAT2 basal extrusion and apical domain constriction **j** schematics of nAT2 extrusion and junctioning outcomes at various interstitial positions.

To further investigate the dynamics of nAT2 junctioning, we performed timelapse confocal microscopy on Precision Cut Lung Slices (PCLS) isolated from e17.5 murine lungs wherein the distal lung epithelium was labeled with a membrane tethered GFP (*Tg^Nkx^*^2^.^1^*^-Cre^ Rosa26^mTmG^*) wherein nAT2s could be readily visualized to extrude and junction (Fig. 2d). We observed both types of nAT2 junctioning - direct when the interstitial distance was short (Fig. 2e) and indirect when greater than a cell length away (Fig. 2f, Suppl. Fig. 2d, Suppl. Video 5). Finally, we observed for most, if not all junctioning nAT2s, a significant reduction in length of MUC1 domains indicative of apical constriction (Fig. 2g, h). Taken together, we believe our observations are of an initial “interlumenal junctioning” event critical in the process of integrating alveolar lumenal surfaces (Fig. 2i, j) that relies upon basal extrusion of nAT2s to junction. Further, we hypothesize the manner of junctioning (direct or indirect) is dependent on interstitial distance, and that defective nAT2 junctioning could result in alveolar simplification similar to what is observed in BPD and COPD.

### Nascent and mature AT2 state differ in fate plasticity and BH3 regulation

Following their initial specification, nAT2s undergo maturation in the first week or so after birth, which can be observed by the switch from RELMα to CD74 expression (Fig. 3a, b). As RELMα expression is lost in mAT2s, we observe its concomitant upregulation in the airway epithelium (Suppl. Fig. 3a). In contrast to the initial proximal-to-distal wave pattern that AT1/AT2 fate selection is reported to occur^4, 13^, we observe AT2 maturation does not follow this pattern but rather occurs in a sporadic salt-and-pepper pattern (Fig. 3c, d, Suppl. Fig. 3b), suggesting maturation may be governed by a distinct mechanism.

**Figure 3:**
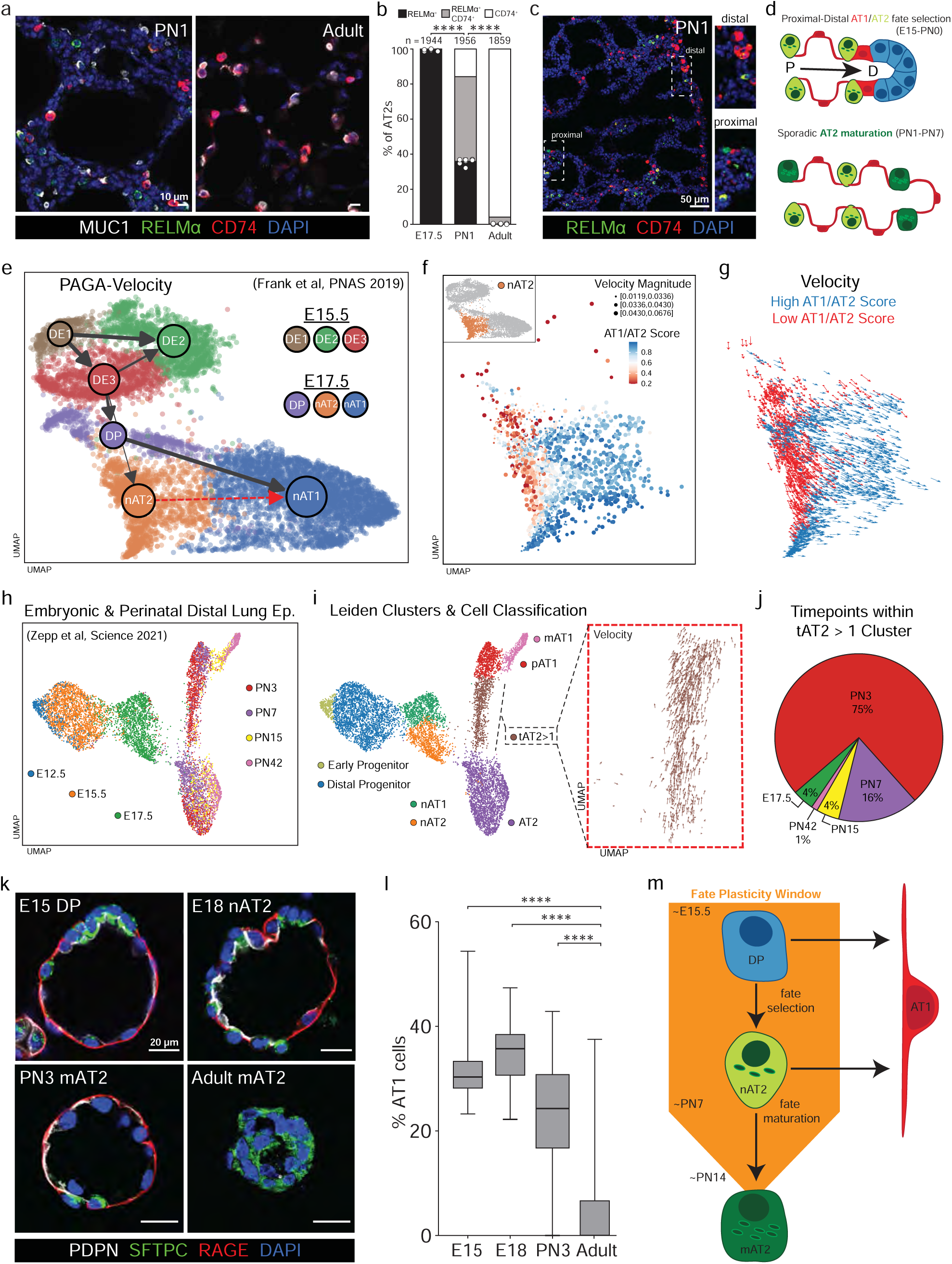
Nascent AT2s retain fate plasticity. **a** PN1 and adult (≥ PN60) lungs immunostained for AT2 marker MUC1 (white), RELMα (green), CD74 (red), and DAPI. Bar, 10 µm. **b** Timecourse quantification of (**a**) showing AT2 transitioning from nascent (RELMα^+^) to mature (CD74^+^). (*n* represents total AT2 cells scored at each timepoint in experimental triplicate, values as mean ± SD). \*\*\*\**p* ≤ 0.0001 (one-way ANOVA, data as mean ± SD). **c** PN1 lung immunostained for RELMα (green), CD74 (red), and DAPI. Close-up showing both CD74^+^ mAT2s and RELMα^+^ nAT2s. Bar, 50 µm. **d** Alveolar fate selection during late embryogenesis, contrasted with AT2 maturation during the early post-natal period. **e** PAGA velocity of scRNAseq data of e15.5 and e17.5 epithelial cells. Arrow thickness represents the relative fraction of cells with velocity from the original cluster to target cluster. A transcriptional trajectory is observed starting from nAT2s to nAT1s (red arrow). **f** Velocity magnitude and AT1/AT2 gene score for the nAT2 fraction from (**e**). **g** Velocity analysis of the nAT2 fraction from (**e**) depicting AT1/AT2 score (cutoff 0.5). **h** scRNAseq data (GSE149563)^37^ of embryonic, perinatal, and adult time points of the distal epithelium. **i** Leiden clustering including a distinct transitional AT2 cluster (tAT2 > 1; brown). Close-up depicts velocity analysis (red dotted box), showing transition towards AT1 cluster. **j** Timepoint distribution of cells observed within tAT2 > 1 cluster in (**i**). **k** Immunostained spheroids of e15.5 distal progenitors (EpCAM^+^), e18.5 nAT2s (RELMα^+^), PN3 mAT2s (MHC-II^+^), and adult mAT2s (MHC-II^+^). Spheroids were cultured on Matrigel for four days with media supplemented every two days with FGF7 (50 ng/mL), followed by fixation and stained for PDPN (white), RAGE (red), DAPI (blue), and AT2 marker SFTPC (green). Cells that remained distal progenitors retained both AT1 markers (RAGE, PDPN) and the AT2 marker (SFTPC). Bars, 20 µm. **l** Quantification of (**k**) showing percentage of cells per spheroid that differentiated into AT1 cells determined by immunostaining (*n* = spheroids sampled for each condition in experimental triplicate). \*\*\*\**p* ≤ 0.0001 (one-way ANOVA). **m** A revised model of the Fate Plasticity Window depicting retained fate plasticity in nAT2 cells with the ability to differentiate into AT1 cells. All experiments were repeated at least three times.

Given the molecular distinction between nAT2 and mAT2 states, we next sought to determine whether functional differences also exist. One manner in which time-dependent cell states can vary is in fate plasticity, as has been observed in neutrophils^35^ in vivo and lung epithelial iPSCs^36^ in culture. To determine whether and how nAT2s and mAT2s differ in fate plasticity, we analyzed two previously generated scRNAseq datasets that sample timepoints either during^12^ (Fig. 3e - g, Suppl. Fig. 3c - g) or far beyond^37^ (Fig. 3h - j) alveolar epithelial differentiation. Analyzing first the distal epithelium prior to (e15.5) and during (e17.5) alveolar epithelial differentiation, we performed standard clustering as well as UMAP visualization and found that distal epithelial cells isolated from these timepoints only rarely co-clustered (Suppl. Fig. 3e), indicating no e15.5 progenitor population had significantly adopted either AT1 or AT2 transcriptional program, a central tenet of the early lineage specification model proposed in a prior study^12^ . A weak correlation between one of the e15.5 progenitor clusters (DE3) and the e17.5 progenitor cluster (0.42) was observed (Suppl. Fig. 3e, yellow box), suggesting a shared transcriptional program.

Single-cell transcriptional dynamics analysis was next analyzed to determine when and how the AT1 and nAT2s clusters segregate. Performing pseudotime (Suppl. Fig. 3f), velocity (Suppl. Fig. 3g), as well as integrated PAGA-Velocity analysis (Fig. 3e) all identified e15.5 clusters as the starting timepoint, as well as identified the AT1 cluster as the predicted endpoint (Suppl. Fig. 3f, g). Further, velocity analysis found a substantial proportion of nAT2s to be on a transcriptional trajectory towards the AT1 cluster, suggesting fate plasticity was retained (Fig. 3e, red arrow). To determine whether a subset of nAT2s displayed a biased trajectory towards AT1 differentiation, we established an AT1 and AT2 score based on well-established marker genes^9^ and observed a clear stratification within the nAT2 cluster (Fig. 3f). Separating the nAT2 cluster by this ratio into two subgroups (High AT1 or Low AT1 gene expression, > 0.5 = High AT1/AT2 score), we compared velocity trajectories and observed a clear separation (Fig. 3g), indicating nAT2 subset with a high AT1/AT2 ratio are leaving the nAT2 state and transitioning to AT1.

After observing that embryonic nAT2s possibly retain fate plasticity, we sought to determine how long AT2s retain fate plasticity after birth. Using a previously reported dataset that samples across the lifetime of the distal epithelium of the mouse lung^37^, we performed similar processing as before to remove duplicates and non-distal epithelial cells before performing UMAP visualization and clustering analysis (Fig. 3h, i). Analyzing more widely over time, we again observed earlier progenitor populations, mature AT1 and AT2 clusters, as well as a cluster transitioning between them as previously reported^37^. Velocity analysis of this intermediate cluster found the cellular trajectories are oriented away from AT2 and towards AT1 (Fig. 3i, red box) like our prior observation. Finally, analysis of the timepoints of cells within the AT2→AT1 transitioning clusters revealed most were from the early perinatal period, suggesting fate plasticity was lost as AT2s mature within the first week after birth (Fig. 3j). To confirm this, we performed alveolosphere differentiation assays following our previously described protocol^10^ and found that both nAT2s as well as early mAT2s (PN3) could give rise to AT1 cells (Fig. 3k, l), confirming fate plasticity was lost primarily in a time-dependent manner, independent of nAT2 or mAT2 state. Taken together, our analysis indicates that fate plasticity, a property of DPs, is retained in newly formed AT2 cells for a period of 1-2 weeks after birth, revising the model of the alveolar epithelial fate selection model to include a window of plasticity (Fig. 3m).

Another property known to vary from early life to maturity is mitochondrial sensitivity to apoptotic stimuli (apoptotic priming)^38^, which is reduced during maturation in several organs but can be retained in certain stem cell populations^39, 40^. To determine whether apoptotic sensitivity is retained in nAT2s, BH3 profiling was performed on isolated DPs, nAT2s, as well as adult mAT2s. As expected, DPs and nAT2s exhibited heightened sensitivity to pro-apoptotic BH3 peptides as indicated by higher rates of cytochrome c release, while mAT2s were largely unresponsive (Suppl. Fig. 3h). Taken together, our findings indicate the nAT2 state retains both fate plasticity as well as sensitivity to BH3 regulated apoptosis despite being functionally differentiated.

### C/EBPα is required to maintain but not select AT2 fate

Given the loss of fate plasticity as nAT2s mature into mAT2s, we next sought to determine how this paradigm is regulated. Referencing the scRNAseq data we previously used to distinguish nAT2 and mAT2 clusters (Fig. 1a, b) as well as AT2s sequenced more deeply by the Tabula Muris Consortium^41^, we screened putative transcriptional and epigenetic regulators from a curated list (1,678 genes) and identified 67 genes with ≥ 2-fold enrichment along the AT2 (Suppl. Table 3) lineage (DP→nAT2→mAT2) compared to AT1 (Fig. 4a). Of these, we then considered only the 32 genes with known embryonic lethal or lung defects when knocked out in mice (Fig. 4a, Suppl. Fig. 4a). These genes can be stratified into four categories based on expression pattern over time: constitutive, DP-restrictive, DP/nAT2-restrictive, and nAT2/mAT2-restrictive (Fig. 4a, lower and Suppl. Fig. 4a). Of these, we focused on genes expressed in the observed window of fate plasticity (Suppl. Fig. 4b), which narrowed the list to four genes, *Etv5* and *Elf5* (constitutive) as well as, *Cebpa* and *Nupr1* (nAT2/mAT2-Restrictive) (Fig. 4b). While both constitutively expressed genes (*Etv5* and *Elf5*) have been investigated in the lung, findings suggest regulatory roles other than fate plasticity, as ETV5 is required primarily to maintain AT2 survival^42^ while ELF5 appears to broadly regulate aspects of lung epithelial differentiation^43^. Further, we excluded study of *Nupr1* given its primary role is in death regulation, not differentiation^44, 45^. This left *Cebpa* as the putative regulator, for which we validated the timing of expression via immunostaining in the embryonic lung (Fig. 4c).

**Figure 4:**
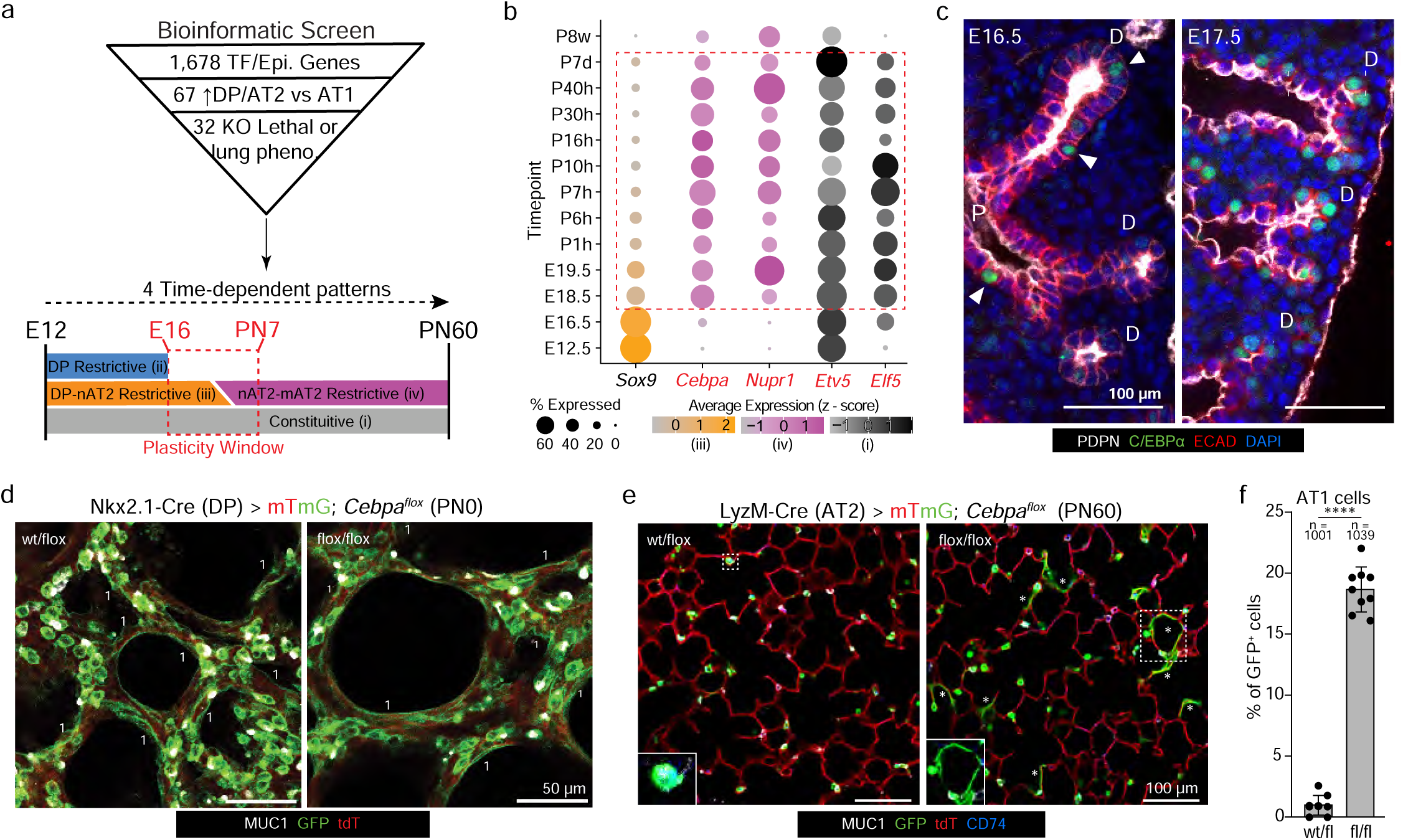
C/EBPα a is required to maintain but not select AT2 fate. **a** A bioinformatic screen of 1,678 transcription factors or epigenetic regulators, out of which 67 are enriched along the AT2 lineage. A final list of 32 genes were selected that have known embryonic lethal or lung defects upon knockout in mice. These 32 genes stratify into four time-dependent categories: DP restrictive, DP-nAT2 restrictive, nAT2-mAT2 restrictive, and constitutive. The Plasticity Window (red dotted box) spans from ∼e16.5 to ∼PN7. **b** Dot plot showing expression of *Sox9* (DP restrictive; yellow), *Cebpa* and *Nupr1* (nAT2-mAT2 restrictive; pink), *Etv5* and *Elf5* (constitutive; grey) over developmental and adult timepoints. The plasticity window is depicted with a red dotted box. **c** e16.5 and e17.5lungs immunostained for PDPN (white), C/EBPα (green), E-cadherin (ECAD, red), and DAPI (blue). C/EBPα^+^ cells are marked (arrowheads) within the proximal (P) and distal (D) regions of the epithelial branches. Bars, 100 µm. **d** Alveolar regions of PN0 control *Nkx2.1^Cre^; Rosa26^mTmG^; Cebpa^wt/fl^* (left) and *Nkx2.1^Cre^; Rosa26^mTmG^; Cebpa^fl/fl^* (right) lungs immunostained for Cre reporter (mTmG) and MUC1 (white). GFP^+^ (DP lineage) cells form both AT1 cells (flattening; “1”) and AT2 cells (MUC1^+^) even upon *Cebpa* deletion. Bars, 50 µm. **e** Alveolar regions of adult control *LyzM^Cre^; Rosa26^mTmG^; Cebpa^wt/fl^* (left) and *LyzM^Cre^; Rosa26^mTmG^; Cebpa^fl/fl^* (right) lungs immunostained for MUC1 (white) and CD74 (blue). AT2-lineage derived GFP^+^ cells are seen to differentiate into flat AT1 cells (asterisk) upon *Cebpa* deletion. Bars, 100 µm. **f** Quantification of (**e**) showing percent GFP^+^ cells that differentiate into AT1s upon *Cebpa* deletion (*n* represents number of GFP^+^ cells sampled for the *Cebpa^wt/fl^* and *Cebpa^fl/fl^* conditions in experimental triplicate). \*\*\*\**p* = 1.4 × 10^-14^ (Student’s two-sided *t*-test, data as mean ± SD). All experiments were repeated at least three times.

Previous *Cebpa* research in the developing mouse lung performed knockouts via a Tet-On approach and concluded it was required for alveolar epithelial differentiation^46, 47^. These findings are conflicting with the expression pattern of *Cebpa*, which is first expressed after AT2 differentiation, not before (Fig. 4b, c, Suppl. Fig. 6b). Given this discrepancy, as well as a subsequent study that questioned the viability of the prior Tet-On approach in the lung^48^, we first sought to confirm whether *Cebpa* was required for alveolar epithelial differentiation via a more refined method of gene deletion. Using a lineage labeling with efficient recombination approach (*Nkx2.1-Cre*; *Rosa26^mTmG/mTmG^*; *Cebpa^fl/fl^*) wherein distal bipotent progenitors are both fluorescently labeled and floxed gene recombined^10^, we tested whether *Cebpa* loss indeed resulted in suppressed differentiation. Surprisingly, we observed that both AT2 and AT1 fate selection occurred despite the absence of *Cebpa* (Fig. 4d, Suppl. Fig. 4c), both at the molecular and morphological level.

As we expected *Cebpa* to play a role later in development consistent to when it is expressed, we next performed sparsely labeling with efficient deletion experiment wherein *Cebpa* was deleted in a subset of AT2 cells starting in the early perinatal period (∼PN5, *LyzM^Cre^; Rosa26^mTmG/mTmG^*; *Cebpa^fl/fl^*). While lineage-traced control AT2 cells maintain their fate into adulthood, a significant portion of AT2 cells lacking *Cebpa* converted into AT1 cells (Fig. 4e, f), suggesting an increase in AT2 fate plasticity. Finally, we confirmed the result using an AT2-restrictive tamoxifen-inducible Cre recombinase line (*Sftpc^CreER^*) which again demonstrated an increase in conversion to AT1 fate both morphologically and molecularly (Suppl. Fig. 4d). Taken together, we determined that C/EBPα, a bZIP transcription factor expressed following AT2 cell specification is required during the perinatal period to maintain, not select, AT2 fate, suggesting it plays a role in suppressing fate plasticity during the transition from nascent to mature AT2 state.

### C/EBPα represses a non-cell autonomous AT1 differentiation program driven by DLK1

To determine more precisely how C/EBPα regulates AT2 fate plasticity, we conducted scRNAseq of the alveolar epithelium from both control (*Cebpa^wt/fl^*) and AT2-floxed (*Cebpa^fl/fl^*) lungs, wherein Cre-mediated deletion was induced in the perinatal period using *Sftpc^CreER^* in combination with the Tg(iSuRe-Cre)^49^ allele. Retrospective identification of cell types via Louvain clustering using Seurat was able to distinguish control (AT2 – *Cebpa^wt/fl^*) and *Cebpa* floxed (AT2 – *Cebpa^fl/fl^*) AT2s as well as AT1s (Fig. 5a-i, ii), a subset of which we confirmed were derived from AT2 cells following Cre induction by expression of the Cre-inducible reporter Tg(iSuRe-Cre) (Suppl. Fig. 5a), confirming our prior lineage tracing experiments. We observed a substantial down- and upregulation of 117 and 116 genes respectively upon loss of *Cebpa* (Suppl. Fig. 5b). To determine how AT2 or AT1 gene expression overall during development (Suppl. Fig. 5c) and was impacted following *Cebpa* loss, we first used gene scores to broadly assess changes in either AT2 or AT1 related genes (Suppl. Fig. 5e). While no significant upregulation of AT1 genes was observed (suggesting C/EBPα likely does not repress fate plasticity by directly repressing the AT1 gene program), a reduction in overall AT2 genes was observed in AT2s lacking *Cebpa* (Fig. 5a-ii). To further determine how AT2-associated genes were altered, we separately assessed genes restrictively enriched in either nAT2 or mAT2 state (Suppl. Fig. 5d). Upon *Cebpa* loss, AT2 cells reduce expression of mAT2 genes while concomitantly upregulating nAT2 gene expression (Fig. 5a-ii), suggesting re-accessing of a plastic, nascent-like AT2 state.

**Figure 5:**
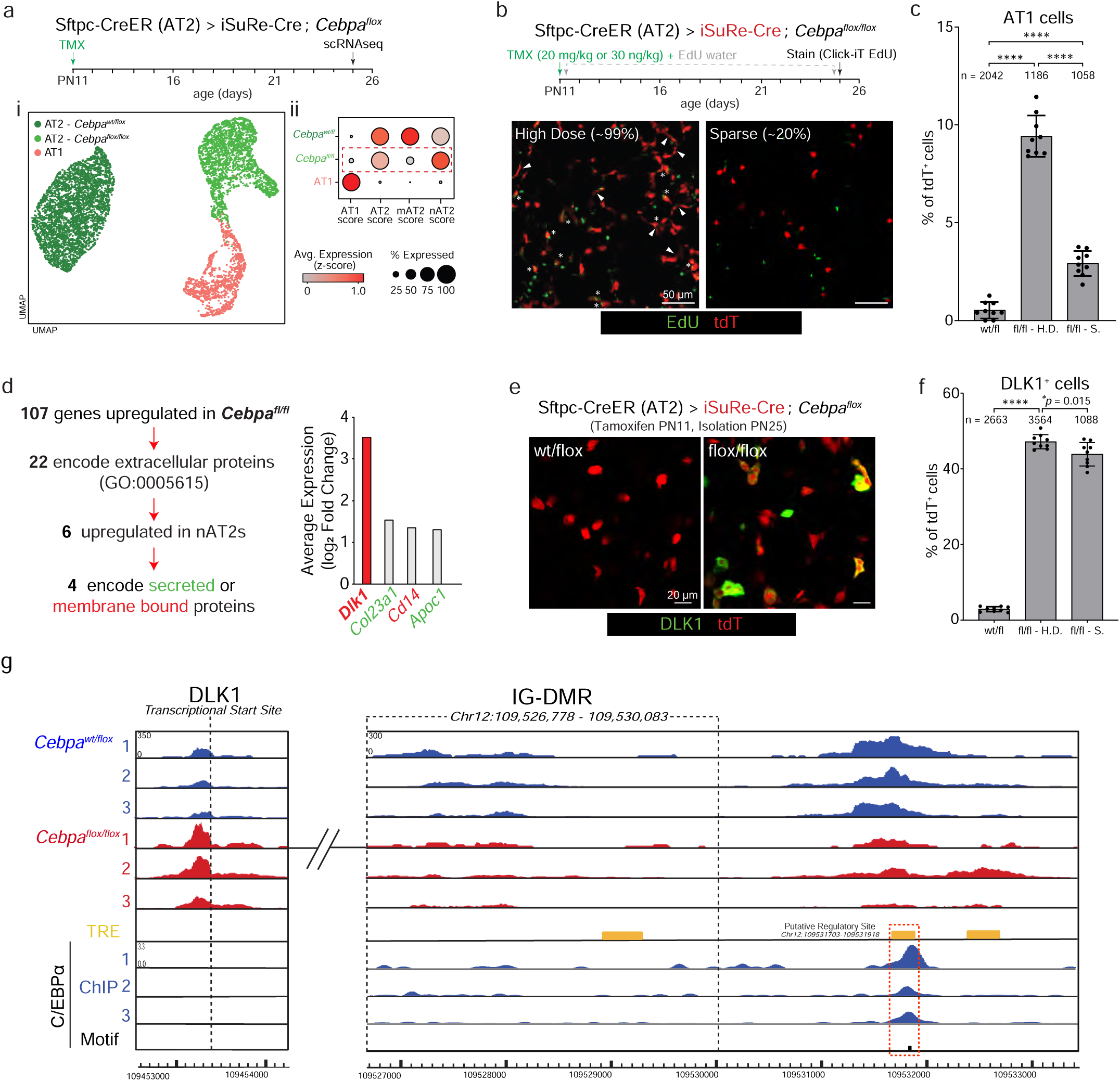
C/EBPα blocks AT2 fate plasticity by repressing Dlk1. **a** Experimental setup and timeline: Lungs were isolated 14 days post TMX (PN25) and alveolar epithelial cells (EpCAM^+^) from control (*Sftpc^CreER^; Tg(iSuRe-Cre); Cebpa^wt/fl^*) and *Cebpa* deleted (*Sftpc^CreER^; Tg(iSuRe-Cre); Cebpa^fl/fl^*) lungs were processed for scRNAseq. **a-i** depicts the clusters (Louvain). **a-ii** shows gene scores across the 3 clusters in (**ai**). Upon *Cebpa* deletion, AT2 and mAT2 scores decreases, while nAT2 score increases compared to control AT2s without activation of AT1 score of the *Cebpa^fl/fl^* cluster (red dashed box). **b** Experimental timeline and TMX dosage. EdU water administered continuously for 14 days. Lungs immunostained for tdT (red) and proliferation (detected by ClickIT in green). *Cebpa* deleted lungs with either high (∼99% AT2s) or sparse dose (∼20% AT2s) TMX, depicting proliferating AT2-lineage cells (asterisk) and flattening AT1 cells (arrowhead). Bars, 50 µm. **c** Quantification of (**b**) showing percent tdT^+^ AT1 cells upon *Cebpa* deletion in either High Dose (fl/fl – H.D.) or Sparse (fl/fl – Sp) deletions compared to control (wt/fl) AT2s (*n* represents tdT^+^ cells sampled for each condition in experimental triplicate). \*\*\*\**p* ≤ 0.0001 (One-way ANOVA with Tukey’s multiple comparisons testing; data as mean ± SD). **d** Average gene expression values (log_2_ fold change) of the 22 (out of 107 total) genes that encode for extracellular proteins enriched in the *Cebpa^fl/fl^* AT2 cluster. Six genes (bold) are found to be upregulated in nAT2 cells and four of which encodes secreted (green) or membrane bound (red) protein, with *Dlk1* (red, bold) having the highest fold expression (log_2_ fold change). **e** Alveolar regions of lungs from control *Sftpc^CreER^; Tg(iSuRe-Cre); Cebpa^wt/fl^*(left) or *Sftpc^CreER^; Tg(iSuRe-Cre); Cebpa^fl/fl^* (right) mice, isolated 14 days post high-dose TMX (at PN25). Lungs were immunostained for DLK1 (green) and tdT (red). Bars, 20 µm. **f** Quantification of (**e**) for percent TdT^+^ AT2 cells that express DLK1 upon loss of *Cebpa* in both high dose and sparse deletion conditions. Note similar percentage of tdT^+^ AT2s express DLK1 in both high dose and sparse deletion conditions, 47.2% and 43.9% respectively (*n* represents tdT^+^ cells sampled for control, high dose and sparse deletion in experimental triplicate). \*\*\*\**p* ≤ 0.0001, \**p* = 0.015 (one-way ANOVA, data as mean ± SD). All experiments were repeated at least three times. **g** Bulk ATACseq tracks showing chromatin accessibility of *Cebpa^wt/fl^*(blue) and *Cebpa^fl/fl^* (red) AT2 cells (*n* = 3 replicates for each condition), showing the *Dlk1* transcriptional start site (TSS) and the intergenic differentially methylated region (IG-DMR). The closest C/EBPα binding site to the *Dlk1* TSS is observed at a distal trans-regulatory element (TRE, yellow), downstream of the IG-DMR, with reduced chromatic accessibility upon loss of *Cebpa*. Further reanalysis of ChIP-seq dataset of the TRE (red dashed box) shows elevated activity at binding sites, indicating their involvement in *Dlk1* regulation.

To better understand the behavior of these plastic AT2s as well as focus our investigation of their molecular regulators, we next sought to determine whether the AT2-to-AT1 differentiation observed upon *Cebpa* loss was mediated by a cell autonomous or non-autonomous mechanism. While a common experimental approach in *Drosophila* research^50, 51^ to delineate between mechanisms targeting internal signaling or proliferation (cell autonomous) versus expression of an extracellular signal (non-cell autonomous), such experiments are less common in the mouse and have been difficult using inducible Cre lines until the development transgenic alleles such as Tg(iSuRe-Cre) ^49^. Usage of a CreER line alone under sparse labeling conditions can result in incomplete or mosaic recombination of alleles and label “escaper cells”^52^, confounding interpretations of the gene’s necessity. To overcome this, We next performed sparse labeling with efficient *Cebpa* deletion using the Tg(iSuRe-Cre) allele (*Sftpc^CreER^; Tg(iSuRe-Cre); Cebpa^fl/fl^*) at a tamoxifen dose we determine labels ∼20% of AT2s validated by immunostaining (Fig. 5b, c, Suppl. Fig. 5f, g). This method of sparse labeling with efficient recombination can be used to delineate whether a phenotype is regulated in a cell autonomous (i.e. via a cell intrinsic mechanism, in which case the phenotype would occur in both sparse and non-sparse labeling experiments) and non-cell autonomous programs (i.e. via a cell extrinsic mechanism, in which case the phenotype would occur significantly less if at all in the sparse labeling experiment). In contrast to when *Cebpa* is removed from most AT2s (∼99%), we observed that removal of *Cebpa* from sparse numbers of AT2s (∼20%) resulted in a substantial reduction of AT2-to-AT1 differentiation as assessed by percentage of lineage labeled AT1s and proliferation (Fig. 5b, Suppl. Fig. 5h). This finding strongly indicates that AT2 fate plasticity acts via a cell non-autonomous mechanism, likely an extracellular signal, to promote AT2-to-AT1 differentiation.

Given this, we focused our screen on genes upregulated in *Cebpa* deleted AT2s that encode proteins with an extracellular localization (GO:0005615), which returned 22 genes, 6 of which were also expressed in the nAT2s from the prior scRNAseq timecourse, and of these 4 are localized on the cell membrane or secreted (*Dlk1*, *Col23a1*, *Cd14*, and *Apoc1*) (Fig. 5d, Suppl. Fig. 5i, j). The most highly upregulated of these genes is *Dlk1*, which encodes a single pass type I membrane protein localized to the outer cell surface that is known to regulate Notch signaling^53^ and AT2-to-AT1 differentiation^23^ (Fig. 5d, red).

Finally, we confirmed DLK1 is also upregulated at the protein level following *Cebpa* loss in culture (Suppl. Fig. 5k), and further observe its expression is maintained under sparse labeling with efficient *Cebpa* deletion in vivo, suggesting C/EBPα directly regulates DLK1 in a cell autonomous manner (Fig. 5e, f). Taken together, we hypothesize that C/EBPα begins to block fate plasticity of specified AT2s by direct suppression of *Dlk1* and its subsequent activation of Notch signaling.

After establishing C/EBPα likely directly represses *Dlk1* expression, we sought to determine where within the genome it is mediated. We performed bulk ATAC-seq on both control and *Cebpa* floxed AT2s in the same experiment timeline as for scRNAseq (Suppl. Fig. 5l), but AT2-lineage cells were further purified by FACS using the genetically encoded fluorophore tdTomato. After confirming correlation between ATACseq replicates (Suppl. Fig. 5m), we identified 23,764 sites within the genome that significantly changed in accessibility following *Cebpa* loss (Suppl. Fig. 5n), with decreased accessibility associated with 3,986 genes, consistent with C/EBPα’s well established role as a positive regulator of gene expression^54^. We observed 369 DEGs shared between our ATACseq and scRNAseq datasets (Suppl. Fig. 5o) and mapped the proportions of genomic features that are up- and downregulated (Suppl. Fig. 5p). We also observed a smaller but substantial number of genes (2,384) with associated upregulation in chromatin accessibility following *Cebpa* loss, indicating repressive regulation, including the transcriptional start site (TSS) of *Dlk1* (Fig. 5g). Further, the change in enrichment of transcription factor binding sites following *Cebpa* loss is consistent with prior work on Nkx2.1 and AP-1 acting as a bivalent co-factor to reinforce fate of both AT1 (via Yap/Taz interactions) and AT2 (via C/EBPα) cells^55^ (Suppl. Fig. 5q, r, s).

In investigating putative regulatory sites near *Dlk1*, we initially considered the nearby intergenic differentially methylated region (IG-DMR) previously demonstrated to strongly regulate *Dlk1* expression^56^. While no change in accessibility was detected in the IG-DMR locus, a region nearby was found to decrease following *Cebpa* loss (Fig. 5g). Further, this region included a predicted regulatory site containing a C/EBPα binding motif (Fig. 5g, dashed red) where it could possibly act to repress *Dlk1* and that we validated C/EBPα binds to in AT2 cells in a previously published ChIPseq dataset^52^.

### PRC2 and C/EBPα act as a *Dlk1* “pulse generator” critical for patterning alveolar epithelial fate

Given the perinatal repression of *Dlk1* by C/EBPα, we were curious of its expression earlier in lung development. As previously reported^57, 58^, we observed that *Dlk1* is lowly expressed at stages prior to alveolar epithelial differentiation (e12.5-e15.5) both at the protein (Fig. 6a) and RNA level (Fig. 6b, dashed grey box). Further, *Dlk1* expression appeared to peak in the period from e17.5 to PN1 (Fig. 6a, b, Suppl. Fig. 6a). As *Cebpa* is not detected at earlier stages (≤ e15.5) (Suppl. Fig. 6b), we were curious how *Dlk1* downregulation is mediated. Suppression of *Dlk1* has also been found to be mediated by the polycomb repressive complex (PRC2), including the H3-K-27 methyltransferase EZH2^59^. We observed that genes encoding PRC2 members were enriched at earlier stages (Fig. 6b, red text), and further that EZH2 protein is substantially downregulated after e16.5 (Fig. 6c).

**Figure 6:**
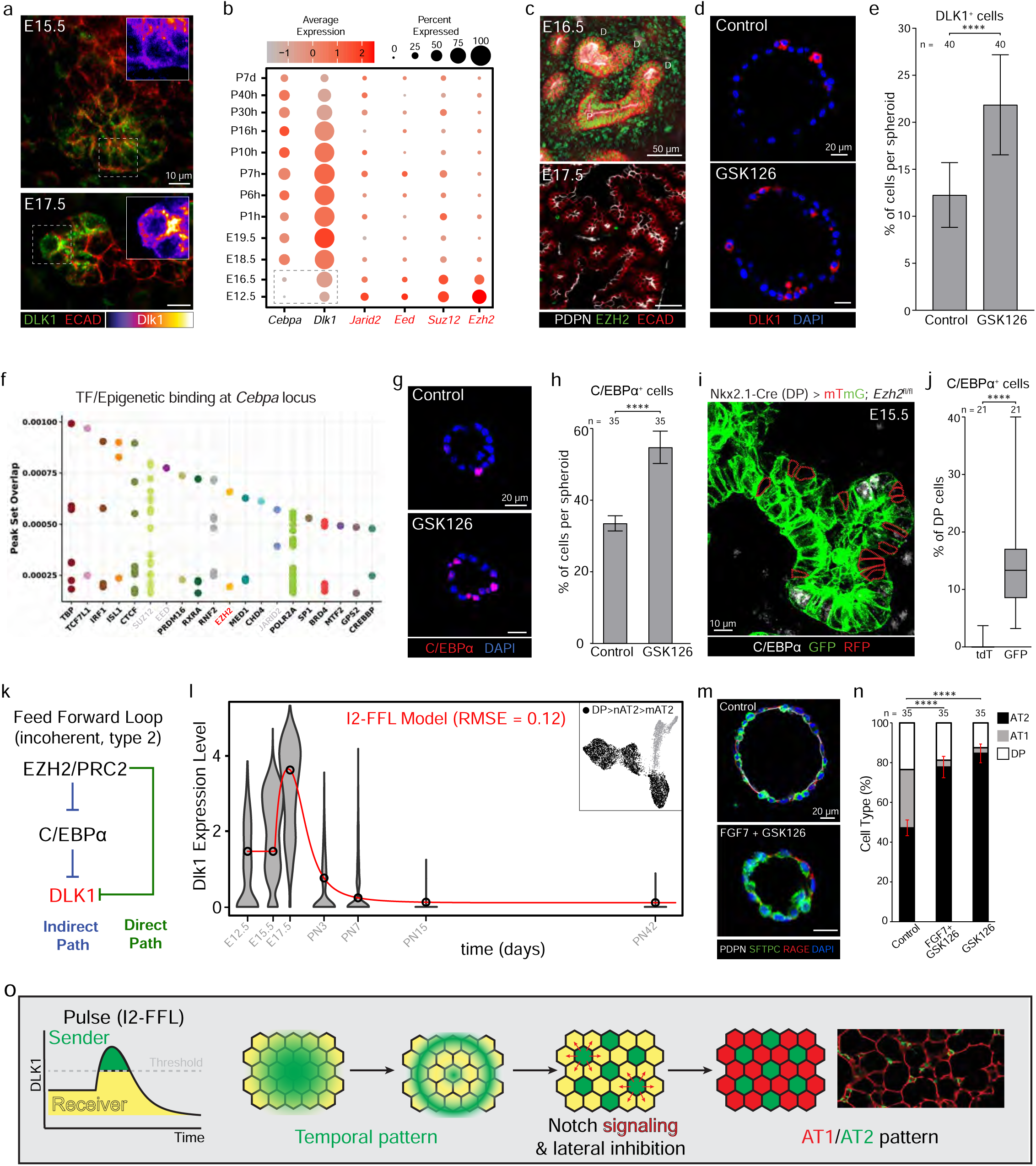
PRC2 and C/EBPα act as an incoherent FFL to generate a DLK1 pulse late in development. **a** Distal alveolar buds of e15.5 (top) and e17.5 (bottom) lungs immunostained for DLK1 (green) and E-cadherin (ECAD, red). Close-up images show e15.5 tips have low DLK1 intensity (fire LUT), and e17.5 tips have higher levels of DLK1 protein. Bars, 10 µm. **b** Dot plot showing RNA expression levels of *Cebpa*, *Dlk1*, and PRC2 members genes (red) during developmental time-course. Minimal expression of *Cebpa* and *Dlk1* observed (grey dashed box), in contrast to the high PRC2 members genes before e16.5. **c** Epithelial branches of e16.5 and e17.5 lungs immunostained for PDPN (white), EZH2 (green), and E-cadherin (ECAD, red). EZH2 presence in the proximal (P) and distal (D) regions of the epithelial branches at e16.5 but absence by e17.5. Bars, 50 µm. **d** Immunostained spheroids of e15.5 DPs (EpCAM^+^) cultured on Matrigel for four days with media supplemented every two days with FGF7 alone (Control, top) or FGF7 with GSK126 (10 µM), an EZH2 inhibitor (bottom). Note an increase in DLK1 (red) in cultures with GSK126. Bars, 20 µm. **e** Quantification of (**d**) showing percent DLK1^+^ cells per spheroid for each condition in the culture at day 4 determined by immunostaining (*n* represents number of spheroids for each condition in experimental triplicate). \*\*\*\**p* = 8.24 × 10^-15^ (Student’s two-sided *t*-test, data as mean ± SD). **f** Cistrome DB Toolkit analysis at the *Cebpa* locus identifies EZH2 (red) and the other PRC2 members (grey) to have enriched binding at this site. **g** Immunostained spheroids of e15.5 DPs cultured as in (**d**). Note an increase in C/EBPα (red) in GSK126 group. Bars, 20 µm. **h** Quantification of (**g**) for percent C/EBPα^+^ cells per spheroid at day 4 determined by immunostaining (*n* represents number of spheroids for each condition in experimental triplicate). \*\*\*\**p* = 3.9 × 10^-36^ (Student’s two-sided *t*-test, data as mean ± SD). **i** Distal epithelial branch of an e15.5 *Nkx2.1^Cre^; Rosa26^mTmG^; Ezh2^fl/fl^* mouse lung, sparsely labeling and deleting *Ezh2* in (GFP^+^) alveolar epithelial cells. Lungs were immunostained for C/EBPα (white) and the Cre reporter (mTmG). Bar, 10 µm. **j** Quantification of (**i**) showing percent of DP cells in the distal alveolar buds expressing C/EBPα in lineage negative (RFP^+^) or lineage positive (GFP^+^) cells (*n* represents number of distal alveolar buds sampled for both groups). \*\*\*\**p* ≤ 0.0001 (Mann-Whitney U-test). **k** Interactions between EZH2/PRC2, C/EBPα, and DLK1 that fits the model of an incoherent, type 2 Feed Forward Loop (I2-FFL). **l** Model fitting of *Dlk1* expression from scRNAseq data of the developing lung^37^, following the DP > nAT2 > mAT2 lineage fitted onto an I2-FFL Model (red line). RMSE = 0.12 (High goodness of fit if RMSE ≤ 0.75). **m** Spheroids of e15.5 DPs cultured as in (**d**), immunostained for PDPN (white), SFTPC (green), RAGE (red), and DAPI. Note that treatment with GSK126 increases the proportion of AT2 differentiation detected by an increase in SFTPC^+^ (green) cells. Bars, 20 µm. **n** Quantification of (**m**) showing a stacked plot of the proportion of DP (white), AT1 (grey) and AT2 (black) cells determined by immunostaining. Compared to control, Both GSK126 alone and FGF7 + GSK126 groups resulted in significantly more AT2 differentiation (*n* represents number of spheroids for each condition in experimental triplicate). \*\*\*\**p* = 3.3 × 10^-43^ for Control vs GSK126, *p* = 2.0 × 10^-36^ for Control vs FGF7 + GSK126 (one-way ANOVA, data as mean ± SD). **o** Schematic of DLK1 pulse upon removal of repressors reopens AT2 fate plasticity window and convert a subset of AT2s into Notch signaling senders. In conjunction of further lateral inhibition, the molecular circuit in (**k**) contributes to AT1/AT2 patterning in lung development. All experiments were repeated at least three times.

As EZH2 and the PRC2 complex are enriched earlier when *Dlk1* is downregulated, we next tested whether EZH2 activity is required to suppress DLK1 in culture. Isolated e15.5 DPs were cultured as organoids and treated either with FGF7 alone or with the EZH2 inhibitor GSK126 (10 µM) for 4 days (Fig. 6d). While control organoids had detectable DLK1 protein (∼12% DLK1^+^ cells), levels significantly increased when treated with the EZH2 inhibitor (∼22%, Fig. 6e), indicating EZH2 methyl transferase activity is required for DLK1 suppression.

When characterizing the expression of *Ezh2* alongside *Cebpa*, we noticed a consistent mutually exclusive pattern (Fig. 6b, Suppl. Fig. 6c), suggesting EZH2/PRC2 might also suppress *Cebpa*. We analyzed a published ENCODE epigenetic dataset that mapped the PRC2 modification H3K27me3 across development^60^ and observed its time-dependent reduction at the *Cebpa* locus concomitant to when *Cebpa* expression is upregulated (Suppl. Fig. 6d). Further, analysis of ChIPseq database of mouse tissues and cells^61^ confirmed significant enrichment in binding of EZH2 as well as other PRC2 components near the *Cebpa* locus (Fig. 6f). To determine whether EZH2/PRC2 is also necessary for repressing C/EBPα, we again conducted organotypic culture with e15.5 DPs during which differentiation was promoted by addition of FGF7 as previously described^10^. Blocking of EZH2 activity by GSK126 significantly increased the percentage of C/EBPα^+^ cells compared to control (55% versus 34% respectively, Fig. 6g, h). Finally, we confirmed whether *Ezh2* is required in vivo to suppress C/EBPα by using a sparse deletion approach wherein a Cre line we previously found to sparsely label DPs^10^ (*Tg^Nkx^*^2^.^1^*^-Cre^*) was bred to a transgenic line to delete *Ezh2* and lineage label by GFP (*Tg^Nkx^*^2^.^1^*^-Cre^ Rosa26^mTmG/mTmG^; Ezh2^fl/fl^*). Immunofluorescence conducted at e15.5, a timepoint where no C/EBPα is observed (Suppl. Fig. 6b), found significant upregulation of C/EBPα in *Ezh2* deleted (GFP^+^) versus control (RFP^+^) DPs residing within the same end buds (Fig. 6i, j).

The interactions between EZH2/PRC2, C/EBPα, and DLK1 we observed appeared to fit the definition of a type 2 incoherent feed-forward loop^62^ (I2-FFL, Fig. 6k). An incoherent feedforward loop (I-FFL) is a type of network motif found in biological systems, such as gene regulatory networks. It contains three components, a regulator, an intermediate , and a target – we propose these components in our model to be PRC2, C/EBPα, and *Dlk1*, respectively (Fig. 6k). The term incoherent means that the regulator (PRC2) acts in a conflicting manner to the target — by both inhibiting (directly on *Dlk1*) and activating it (indirectly through repression of the repressive intermediate C/EBPα. One emergent property of an I-FFL is the capacity to generate a pulse of the target gene’s expression (*Dlk1*) following downregulation of the upstream regulator (EZH2/PRC2). To more precisely assess whether the observed temporal pattern of *Dlk1* expression matched the pulse predicted by and I2-FFL, we perform modeling using the biocircuits package (https://biocircuits.github.io/). After establishing reasonable values for parameters describing the relationship of factors in the I2-FFL (see Methods), the model was found to fit well to the observed *Dlk1* expression pattern as assessed by root mean squared error (RMSE = 0.12, Fig. 6l, Suppl. Fig. 6e). Next, we wanted to assess the importance of this molecular circuit in patterning alveolar epithelial differentiation. Immunostaining of e15.5 DPs in organotypic culture for markers of AT2 and AT1 fate (SFTPC and RAGE, respectively) revealed that blocking EZH2 activity, with or without FGF7 treatment, significantly increased the proportion of molecularly specified AT2s while concomitantly reducing AT1s compared to FGF7 alone (Fig. 6m, n). Finally, we wanted to assess whether lateral inhibition of Notch signaling occurs in AT1 cells, as it has been well demonstrated to promote a spatial “salt and pepper” pattern of cell fates^63^. Analysis of scRNAseq timecourse datasets found that nascent AT1 cells upregulate the Notch target gene *Hes1* suggesting signaling upregulation as expected from prior studies^64^ as well as downregulation of Notch ligands *Jag1* and *Dll4* that is consistent with lateral inhibition of Notch signaling (Suppl. Fig. 6f, g). Further, we also observe in nAT1s changes in Notch regulators that is also consistent with lateral inhibition, specifically downregulation of *Dlk1* and upregulation of *Dlk2* (Suppl. Fig. 6f, g) - the latter has been found to act as a cis-inhibitor of DLK1^65^. Therefore, our findings suggest that the potential model of *Cebpa*-mediated fate regulation is as follows (Fig. 6o): In the *Cebpa^fl/fl^* background, the AT2 fate plasticity window is reopened, and the pulsing of DLK1 upon the removal of repressors converts a subset of AT2s into Notch signaling sending cells. Through further lateral inhibition, the molecular circuit depicted in Fig. 6k likely contributes to AT1/AT2 patterning during lung development. Taken together, our findings support the model that EZH2/PRC and C/EBPα act as a molecular circuit (specifically an I2-FFL) to generate a DLK1 pulse during late-stage embryogenesis that is critical to patterning the proportion of AT2 and AT1 cells.

### C/EBPα suppression is required for accessing both reparative and defensive AT2 states

Given the role of C/EBPα in suppressing plasticity in AT2 cells, we sought to determine if and how it is overridden following injury in the adult lung, where AT2s are well established to act as facultative progenitors to regenerate lost AT1 cells^4, 22^. To first determine if C/EBPα downregulation is required following injury in adulthood for AT2-to-AT1 conversion, we infected adult mouse lungs harboring a Cre-dependent fluorescent reporter (*Rosa26^mTmG/mTmG^*) with AAVs targeting AT2 cell expression via a minimal *Sftpc* promoter as previously described^10^ to express either Cre alone or Cre and C/EBPα. After five days, we injured lungs via intratracheal instillation of the irreversible FGFR inhibition Fiin1 (14.75 mg/kg, as previously characterized by our group^10^) or intraperitoneal injection of BHT (225 mg/kg) and performed immunostaining 14 days afterwards (Fig. 7a upper, Suppl. Fig. 7a, b). Intratracheal instillation of AAVs efficiently labeled AT2s with high specificity (Suppl. Fig. 7c). While injured lungs showed AT2-to-AT1 conversion when treated with the control AAV, it was significantly reduced in those given C/EBPα overexpressing AAVs (Fig. 7a, b).

**Figure 7:**
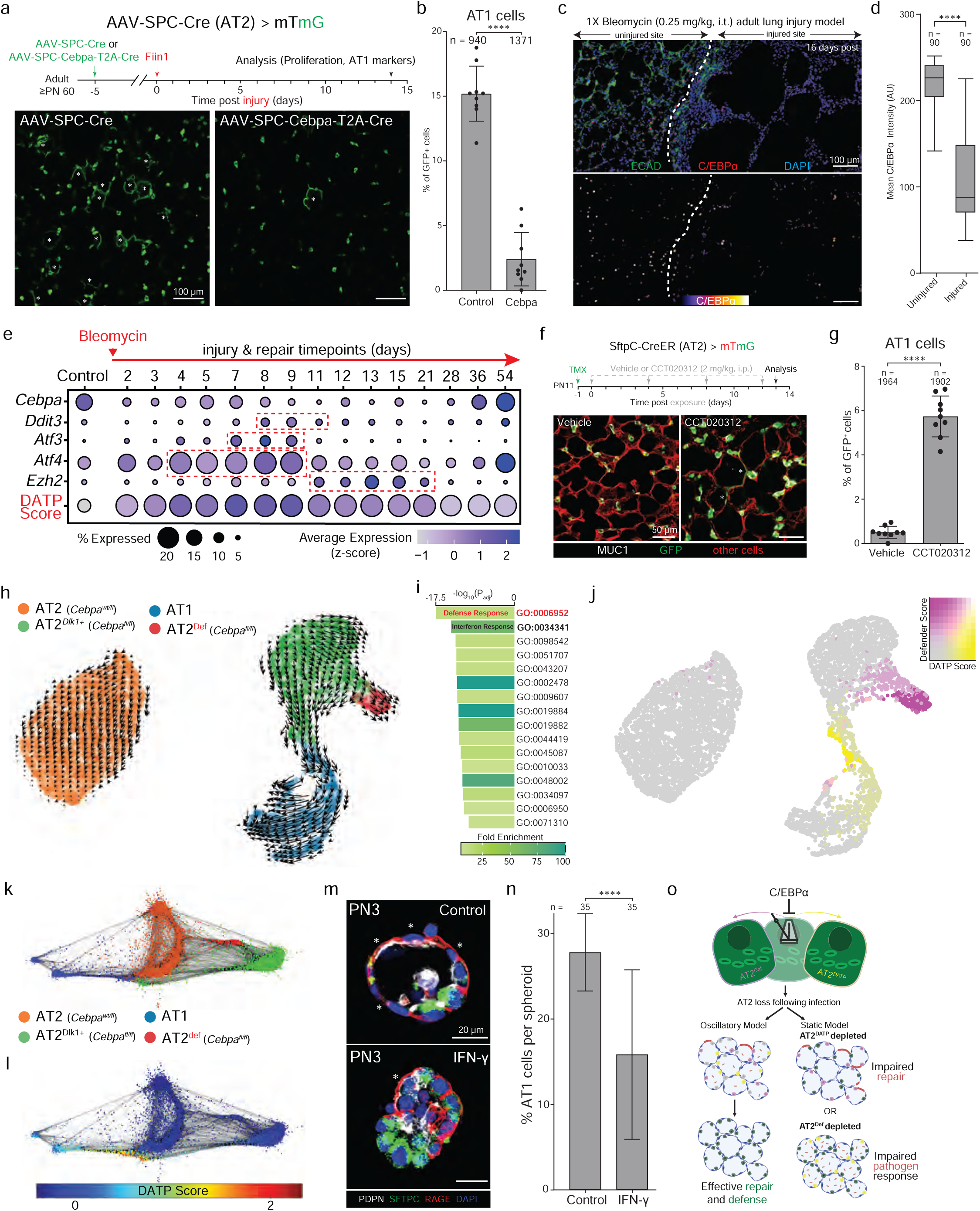
C/EBPα downregulation is required to re-access mutually exclusive Defender and Regenerative states in response to injury. **a** Experimental design (top). 5 days after AAV-SPC-Cre (Control; left) or AAV-SPC-Cebpa-T2A-Cre (right) instillation into lungs of adult *Rosa26^mTmG^* mice to induce constitutive GFP-Cre expression and overdrive expression of *Cebpa* in AT2 cells, lungs were injured by Fiin1 (14.75 mg/kg) intratracheal instillation (i.t.). Lungs were isolated and immunostained for Cre reporter (GFP) and AT1 markers 14 days post injury (dpi). Immunostained images (bottom) show GFP^+^ AT1 cells (asterisk) in the control, but little to none when C/EBPα is overexpressed. Bars, 100 µm. **b** Quantification of (**a**) for percent GFP^+^ cells that differentiated into AT1 cells (*n* represents GFP^+^ cells sampled for each condition in experimental triplicate). \*\*\*\**p* = 6.5 × 10^-10^ (Student’s two-sided *t*-test, data as mean ± SD). **c** Alveolar regions of an adult mouse lung 16 days post 1X Bleomycin (0.25 mg/kg, i.t.) injury, immunostained for E-cadherin (ECAD, green), C/EBPα (red), and DAPI (blue). Injured (right) sites show simplified alveolar structures with reduced E-cadherin expression compared to uninjured sites (left). C/EBPα breakout (fire LUT) below depicts reduced C/EBPα expression in injured versus uninjured sites. Bars, 100 µm. **d** Quantification of (**c**) for mean C/EBPα intensity in AT2 cells within uninjured or injured regions (*n* represents cells scored for each condition in experimental triplicate). \*\*\*\**p* ≤ 0.0001 (Mann- Whitney U-test). **e** Dot plot from a scRNAseq dataset (GSE141259)^20^ of 1X bleomycin-induced injury in AT2 cells, showing relative gene expression in vehicle (control) and various timepoints post 1X Bleomycin injury. Spikes in expression of DATP state associated genes (*Ddit3*, *Atf3*, *Atf4*, and *Ezh2*) are marked (dashed red box), whereas a temporary reduction of *Cebpa* is observed upon injury. The DATP gene score (red, see Suppl. Table 1) is also plotted on this timecourse, showing an activation of a DATP associated state upon injury, which resolves over time. **f** Experimental design for lineage labeling AT2 cells at PN11 in *Sftpc^CreER^; Rosa26^mTmG^* mice, followed by administration of corn oil (vehicle) or CCT020312 (2 mg/kg, intraperitoneal), a Ddit3 agonist, 1 day post TMX, and every 4 days (4 doses total). Lungs were isolated 14 days post TMX (PN25) and immunostained for Cre reported (mTmG) and MUC1 (white). Upon treatment with CCT020312, an increase in GFP^+^ AT1 cells (asterisk) is observed. Bars, 50 µm. **g** Quantification of (**f**) showing percent GFP^+^ AT1 cells for either vehicle or CCT020312 conditions (*n* represents GFP^+^ cells sampled for each condition). \*\*\*\**p* = 2.1 × 10^-11^ (Student’s two-sided *t*-test, data as mean ± SD). **h** Leiden clustering of our scRNAseq dataset reveals two distinct *Cebpa^fl/fl^*AT2 clusters: AT2^DLK1+^ (green) and AT2^Def^ (red). Velocity analysis (arrows) show AT2^DLK1+^ cells going in two separate trajectories, towards the AT1 and AT2^Def^ clusters. **i** Gene Ontology (GO) analysis of the AT2^Def^ cluster shows enriched pathogen defense response (red) and interferon signaling (black) gene programs. **j** DATP- and Defender-associated gene score co-expression shows a mutually exclusive pattern in *Cebpa^fl/fl^* cells. **k** SPRING plot of the four clusters observed in (**h**), showing AT2^Def^ state (red) in between the control AT2 (orange) and AT2^DLK1+^ (green) clusters. **l** SPRING plot of the DATP-associated gene score showing a high DATP score between the control AT2 (middle) and AT1 (left) clusters. **m** Immunostained spheroids of PN3 mAT2s cultured on Matrigel for four days with media supplemented every two days with FGF7 alone (Control, top) or with interferon-γ (20 ng/mL) (bottom). Note a decrease in AT1 cells (asterisk) in cultures treated with interferon-γ. Bars, 20 µm. **n** Quantification of (**m**) showing percent of cells per spheroid that differentiated into AT1s for each condition (*n* represents spheroids sampled per condition). \*\*\*\**p* = 1.2 × 10^-8^ (Student’s two-sided *t*-test, data as mean ± SD). **o** Schematic showing AT2 oscillation between AT2^Def^ (purple) and AT2^DATP^ (yellow) states, negatively regulated by C/EBPα. Upon AT2 loss (grey AT2s) due to infection, remaining AT2s can oscillate (left) between the two states resulting in proper infection response and regeneration. The inability to access either the AT2^Def^ or AT2^DATP^ states would result in impaired pathogen response or impaired repair and regeneration respectively. All experiments were repeated at least three times.

Next, we examined whether C/EBPα expression is downregulated following injury using the 1X Bleomycin model, which results in patchy regions of injury and that resolve within 4-6 weeks. Bleomycin treated lungs were imaged 16 days after administration and injured areas detected by alveolar simplification and reduction in E-cadherin as previously observed^66^.

Immunostaining for C/EBPα revealed a significant downregulation in injured versus uninjured regions (Fig. 7c, d). This finding is in alignment with analysis of a recent scRNAseq timecourse study^20^ following single-dose bleomycin, during which *Cebpa* expression is downregulated in AT2 cells as early as 2 days post bleomycin (Fig. 7e).

After establishing the presence and necessity of C/EBPα downregulation following injury, bioinformatic screening of the bleomycin injury timecourse for transcriptional regulators known to suppress C/EBPα that are also upregulated in the previously identified AT2-to-AT1 transitionary^20^ state (Krt8^+^ ADI, also known as DATP^32^ or PATS^67^) identified three genes, *Atf3, Atf4,* and *Ddit3* (Fig. 7e, Suppl. Fig. 7e). Both activating transcription factors 3 and 4 are known to bind the *Ddit3* promoter to drive expression^68, 69^, and DNA damage-inducible transcript 3 protein, also known as C/EBP homologous protein (or CHOP), has been well established to directly bind and act as a dominant negative inhibitor of C/EBP transcription factors^70^. Given its direct interaction and inhibition of C/EBPα, we next wanted to determine whether induction of CHOP is alone sufficient to drive AT2-AT1 conversion. A day after lineage labeling AT2 cells in lungs of perinatal mice (PN11, *Sftpc^CreER^; Rosa26^mTmG^*), intraperitoneal injections of either vehicle alone or a small molecule inducer of CHOP via PERK activation CCT020312 (2 mg/kg) were administered every four days and lungs isolated at PN25 (Fig. 7f). While little to no lineage labeled AT1s were observed in vehicle treated lungs, there was a significant increase in those treated with CCT020312 (Fig. 7f, g), suggesting PERK/CHOP induction can promote AT2-to-AT1 conversion.

To precisely map AT2 transcriptional trajectories and programs following *Cebpa* loss, we conducted Velocity analysis as well as Leiden clustering on our scRNAseq dataset and confirmed a cellular trajectory from AT2 to AT1 cluster as expected (Fig. 7h). Unexpectedly, we observed another distinct trajectory terminating on a distinct AT2 cell cluster that was enriched in genes associated with pathogen defense response and interferon signaling by GO analysis (Fig. 7i, Suppl. Table 2), leading us to name the “defender” state AT2^def^ (Fig. 7h, red).

Visualizing the expression of AT2^def^ enriched genes versus those associated with the damage-associated transitionary progenitor (DATP) state^32^, we observed these programs were largely mutually exclusive, apparent by both UMAP (Fig. 7j) and SPRING plot (Fig. 7k, l). Given the distinct trajectories, as well as prior research indicating interferon signaling negatively impacts lung epithelial repair^24^, we were interested whether stimulation of the AT2^def^ state by interferon signaling would block AT1 conversion in culture. Given the expression pattern of interferon receptors as well as transcriptional targets in the AT2^def^ cluster (Suppl. Fig. 7d, e), we treated cultures with interferon-γ (Fig. 7m). While perinatal AT2s could give rise to AT1s following FGF7 treatment in organotypic culture, addition of exogenous interferon-γ resulted in a significant reduction AT1 conversion, consistent with the model that AT2^def^ and AT2-to-AT1 molecular program are antagonistic to each other at the cellular level (Fig. 7m, n). To assess the degree to which these programs are present in the same or separate AT2 cells following injury, we analyzed scRNAseq datasets wherein lungs were injured in distinct ways, including diphtheria toxin ablation of AT1 cells^71^, influenza^72^, LPS^73^, and Bleomycin^20^. In support of our findings following *Cebpa* deletion, DATP and Defender states were found to primarily exist in separate AT2 cells during the repair process across all forms of injury (Suppl. Fig. 7f).

## Discussion

These results show that AT2 fate plasticity exists in a time-window from late embryonic to early postnatal life consistent in time with nascency. The timing and length of this window is dictated by a molecular circuit driven by PRC2 and C/EBPα that together generate a pulse of *Dlk1,* a known regulator of adult AT2 fate plasticity. Downregulation of C/EBPα occurs following injury in the adult lung, is required to regenerate AT1 cells, and is mediated in part by the dominant negative C/EBP family member CHOP. C/EBPα also suppresses an AT2 defensive state that is mutually exclusive from fate plasticity that appears to be driven by interferon signaling.

From characterizing its nascent state, we determined AT2 cells first emerge around e15.5 at intermediate regions between the tips and stalks of the distal branches. Our findings consolidate differing models on AT2 emergence, with our observed timing supporting the bipotent model^4, 9, 10, 33, 37^ and our observed pattern suggesting an intermediate starting point, distinct from that proposed in both the bipotent (more proximal) and earlier differentiation^12^ (distal tip) models. After initial emergence (e15.5-16.5), subsequent AT2s next arise at proximal regions coinciding with ASM-to-myofibroblast differentiation as previously reported^34^. Taken together, these findings suggest more than one cue may be present to suppress alveolar epithelial differentiation, one distal and another proximal in position. While it has been established that distal tips are enriched in Wnt activity and thus retain stemness^74^, it is unclear how more proximally positioned DPs are suppressed. One compelling possibility is that ASM directly suppresses differentiation of these proximally localized distal progenitors through direct interaction on the basal epithelial surface, although it is unclear how said suppression is molecularly mediated.

Our findings on AT2 emergence have also provided insight into the sequence of their morphological changes. Given that basal extrusion occurs after AT2 molecular specification, it is likely downstream of fate selection rather than an upstream regulator as previously proposed^75^.

This observation is supported by prior work wherein AT2 fate selection was found to occur under culture conditions that block basal extrusion^10^. If a byproduct of fate selection, what purpose does basal extrusion serve? Following extrusion, we observe that some extruded AT2 cells engage with nearby epithelium from anatomically distinct branches, forming what we name interlumenal junctions, or ILJs. We hypothesize basal extrusions are required to form these junctions, that they are mediated in part by contraction of the AT2 lumenal surface, and likely play a critical early role in postnatal alveologenesis, the stitching together of alveolar lumens^76^. While this process has been observed at the tissue level^14^, as has the multilumenal nature of mature AT2s^34^, many questions remain as to the underlying cellular and molecular mechanisms governing it. Further work on interlumenal junctions will be critical to understanding this stage of alveolar maturation and assessing whether its disruption could underly diseases such as BPD and COPD.

Following their emergence, nascent AT2 cells undergo transition towards their mature AT2 transcriptional state over the course of ∼7 days. This process occurs in a sporadic, salt and pepper pattern distinct from the reported proximal-to-distal wave of alveolar epithelial fate selection^13, 77^ suggesting the cues driving maturation might arise from a different source, which is likely to be immune given the genes enriched in mAT2s as well as prior studies demonstrating interactions underlying one another’s maturation^33^ and function^15^. While AT2s have been shown to impact macrophage maturation^33^, an immune cell type has not been implicated in driving AT2 maturation during postnatal development. More work is needed to better understand which immune cell types interact with AT2 cells during this process and determine how they impact differentiation. While AT2 cells from both nascent and mature states retained plasticity, mAT2s quickly lose this capability within the first week or so after birth, suggesting fate plasticity is somewhat negatively correlated to AT2 maturation. Given these findings, we revised our model of alveolar epithelial differentiation to include a fate plasticity window extending from e15.5 distal progenitors into AT2 nascency, and that tapers down over 1-2 weeks after birth. It is likely this window is critical for postnatal alveologenesis and might be involved in the observed age-dependent severity of COVID19^78^.

While our bioinformatic screening for transcriptional regulators of the plasticity window identified C/EBPα as a possible repressor, previous genetic deletions studies concluded that *Cebpa* is required earlier for both AT1 and AT2 differentiation^46, 47^. As these findings conflict with the observed expression pattern of C/EBPα, which is upregulated following AT2 fate selection, we conducted sparse *Cebpa* deletion experiments in both DP and AT2 populations.

From these studies we found that *Cebpa* is dispensable for the selection but required for the maintenance of AT2 fate, with the later finding recently confirmed in a manuscript by another group^52^. The discrepancy is likely the result of the field’s prior use of genetic tools now found to cause toxicity in the developing mouse lung^48^. While our subsequent scRNAseq and ATACseq analysis reveals repressive targets of C/EBPα, its positive transcriptional regulation of gene expression has been demonstrated including recent publications ^52^. While *Cebpa* loss does result in a reduced expression of several AT2 mature genes, identified in both studies, it is important to note that these AT2 cells do not concomitantly increase the expression of AT1 genes nor directly convert into AT1 cells. Rather the AT2 program reverts to the nascent AT2 state, again suggesting a repressive mechanism is at play that targets fate plasticity. While Sox9 was reported in the other study to be upregulated following *Cebpa* loss, we only detected a modest increase at the RNA level and screening of scRNAseq data referencing a DP gene list revealed no strong upregulation, supporting our model that loss resulted in reversion to a more nAT2 state rather than full dedifferentiation back to DP. Recent work has found that AT1s in the perinatal period also retain fate plasticity and when stretch and YAP/TAZ signaling is lost they are capable to AT1-to-AT2 differentiation^79^. While more work is needed to integrate those findings with ours and others^52^, one compelling model is that both perinatal AT2 and AT1 cells retain fate plasticity for a period of time but are each suppressed by distinct repressors – C/EBPα for AT2s and YAP/TAZ for AT1s.

Our investigation into the key gene(s) repressed by C/EBPα that could regulate fate plasticity identified *Dlk1*, which encodes a cis-regulator of Notch signaling previously implicated in AT2-to-AT1 conversion^23^. Notch is distinct from other signaling pathways in its use of cis-regulators (like DLK1) for activation and has been shown to be critical in generating its well characterized “salt and pepper” pattern^80^. While regulation of the *Dlk1* locus is well documented to be mediated by an intergenic differentially methylation region (IG-DMR) downstream of the gene^56, 81^, ATAC-seq revealed no change in its accessibility nor any changes immediately upstream of the gene’s transcriptional start site upon loss of *Cebpa*. Rather, a dramatic change in accessibility was observed in a previously uncharacterized region roughly 2 kilobases downstream of the IG-DMR that the UCSC genome browser predicts is an enhancer. More work is needed to determine how this element is regulated during development as well as repair following injury in the adult lung.

Our investigation into developmental regulation of *Dlk1* expression determined that PRC2 and C/EBPα act as a type 2 incoherent feedback loop to generate of pulse of DLK1 following downregulation of the PRC2 complex and likely its repressive epigenetic mark, H3K27me3. While several feedback loops have been postulated and constructed in the field of systems biology^62^, most of the loops observed in eukaryotic cells are either type 1 coherent or incoherent in configuration^62^. While the identified loop type (I2-FFL) is observed less frequently, we believe the PRC2-C/EBPα regulation plays a critical role in timing and pattern of alveolar epithelial differentiation, and future studies are needed to test whether reactivation of said loop could be used to re-access AT2 fate plasticity in diseased lungs. We believe this feedback loop constitutes a novel molecular circuit – a “pulse generator” of Notch activation, resulting in spatial patterning of AT1 and AT2 differentiation through lateral inhibition. It is also conceivable that disruption of this molecular circuit could result in an aberrant AT2 state that is disease associated, such as those observed in models of IPF^82, 83^.

Finally, our investigation into the adult lung demonstrates that C/EBPα downregulation following injury is required for repair and is partly mediated by a C/EBP family member with dominant negative activity, CHOP^70^. As CHOP is upregulated from various types of injury that cause ER stress^84^, it is still unclear how C/EBPα downregulation is stimulated following injury of alveolar epithelial cell death, especially as we observe little to no C/EBPα downregulation upon sparse AT2 ablation (Suppl. Fig. 7e). One compelling model requiring further inquiry is that AT1 death sends a specific cue that downregulates C/EBPα to promote fate plasticity in nearby AT2 cells and thus alveolar repair. Interestingly we found that *Cebpa* loss also allowed AT2 cells to access what appears to be a pathogen defensive state driven by interferon signaling. While presence of such a state is expected, as AT2 cells have been implicated in playing a role in immune response in the lung^15^, said state was not expected to be suppressed by C/EBPα nor to be a mutually exclusive transcriptional state from the well-established regenerative DATP state^67^,

Taken together, these observations suggest that C/EBPα might suppress a larger program that alternates between two states – one regenerative and the other defensive. If true, said program would represent a more robust response to pathogen-mediated injury then previously appreciated, as alternating between regenerative and defense states could ensure enough AT2s are available at a given time to both promote clearance of the pathogen as well as repair damaged alveoli (Fig. 7o).

We will briefly cover the limitations of this study to motivate and direct future work. First, we dissociate murine lungs to form organoids which, while a common approach in the field^10, 22, 34^, might alter cellular phenotypes and promote fate plasticity. While the observed behaviors of cultured DPs and AT2s largely match in vivo experiments^10, 22, 34^ and velocity analysis (Figure 3e-j), future work investigating fate plasticity more precisely in vivo would be informative. Differences observed by normalized gene expression alone would benefit from validation via ISH or immunostaining. As our study does not investigate whether AT1s derived from DPs are distinct from those from AT2s, more work is needed to make said delineation. As conclusions of scRNAseq cluster similarity can be impacted by cut-off points, more work delineating differences in e15.5 DPs in an unbiased manner would be informative. While we focused on the role of C/EBPα, future work on other genes from our screen would likely identify other critical regulators of AT1/2 fate plasticity (see Suppl. Table 3). As BH3 death regulation is studied only to determine its similarity between DPs, nAT2s, and mAT2s, more research is needed to determine its role during development and repair. As CHOP expression was induced using a smaller activator of PERK (CCT020312) that may stimulate off target molecules conflating our experiments, further work is needed to precisely investigate the function of CHOP in vivo. Finally, as our experiment revealing C/EBPα’s regulation of a defensive AT2 state did not directly introduce a pathogen, future work is needed to confirm said state is upregulated following infection.

## Methods

### Ethics

All mouse experiments followed applicable regulations and guidelines and were approved by the Institutional Animal Care and Use Committee at the Mayo Clinic (Protocol# A00006319-21).

Mice were house under a 12 h light/12 h dark cycle and lights were not used during the dark cycle. Mouse housing temperature was kept at 65-75 °F (∼18-23 °C) and humidity at 40–60%. Mice were housed in filtered cages and all experiments were performed in accordance with approved Institutional Animal Care and Use Committee protocols and ethical considerations at the Mayo Clinic.

### Mouse strains

Timed-pregnant C57BL/6 J females (abbreviated B6; Jackson Laboratories) were used for all embryonic time points, with gestational age verified by crown–rump length. For studies of adult wild-type lungs, B6 males and females were used. Sparse labeling and recombination studies were conducted by Cre recombinase expression using gene-targeted alleles BAC-*Nkx2.1^Cre^*, LyzM (also called Lyz2)-Cre, and *Sftpc^Cre-ERT^*^2^*^-rtTA^*^58^. Cre-dependent target genes were the conditional “floxed” *Cebpa*^86^ and *Ezh2*^87^ for removal of *Cebpa* and *Ezh2* respectively. Two Cre reporters were used: *Rosa26^mTmG^* ^88^ (JAX stock #037456) and Tg(iSuRe-Cre) (MGI:6361145) ^49^. Upon recombination, the Tg(iSuRe-Cre) allele labels the cell with membrane and cytoplasmic tdTomato (tdT) and constitutively expresses Cre recombinase in the targeted cell, thus ensuring that all Cre-conditional alleles are recombined, even under sparse or clonal conditions. For initiating recombination with *Sftpc^Cre-ERT^*^2^ (*Sftpc^CreER^*), intraperitoneal injection of tamoxifen was administered either at 20 mg/kg for complete (high dose) recombination in adult mice (> PN60), or at 30 ng/kg for sparse recombination in perinatal mice (P0-P21), except where noted. EdU (Cayman Chemicals 20518) was administered via drinking water at 0.1 mg/mL for the indicated period of time. Genomic DNA was extracted from ear punches followed by Proteinase K (KAPA Biosystems) digestion, and genotyping was performed by polymerase chain reaction (PCR) using published primer sets.

### Single-cell RNA-seq bioinformatic analysis

#### Cell Isolation for library preparation

Cells from PN25 *Sftpc^CreER^; Tg(iSuRe-Cre);Cebpa^wt/fl^*and *Sftpc^CreER^; Tg(iSuRe-Cre);Cebpa^fl/fl^* mice (2 weeks after TMX) were isolated and enriched (EpCAM^+^) as per protocol described below.

#### scRNAseq library preparation and processing

Single cell RNA sequencing was performed using the Chromium Single Cell 3’ system (10x Genomics) at the single cell sequencing core at Mayo Clinic according to manufacturer’s instruction (10x Genomics). Sequencing depth is an average of 75k reads per cell to detect lowly expressed genes. Fastq files were generated using Cellranger v 7.2. The sequenced files were mapped to the GRCm38/mm10, supplemented with tdTomato, PhiMut, and WPRE transcripts as per the Tg(iSuRe-Cre) construct^49^.

#### scRNAseq data analysis

We used Scanpy^89^ and Seurat v.4.2.3^90^ for processing and downstream analysis. In the Scanpy pipeline, we identified highly variable genes using the scanpy function scanpy.pp.highly_variable_genes with arguments min_mean = 0.0125, max_mean = 3, and min_disp = 0.5. We restricted to this set of genes for downstream analysis. We further used scanpy.pp.regress_out to regress out unwanted variation in the count of Unique Molecular Identifiers (UMIs) and the percentage of mitochondrial reads. We ran scanpy’s built in Principle Component Analysis (PCA) algorithm with sc.tl.pca and default parameters. We then created a nearest neighbor graph using sc.pp.neighbors with 200 neighbors and 50 Principle Components. We then produced a UMAP dimensionality reduction with scanpy.tl.umap using min_dist = 0.001. We identified cell clusters through unsupervised clustering using the Leiden clustering algorithm through with resolution = 0.8. We assigned cell labels to the clusters based on marker gene expression, and removed several contaminating mesenchymal clusters. These clusters were then visualized on the UMAP.

In the Seurat pipeline, doublets were estimated and removed using the DoubletFinder Package^91^. Dead cells or outliers were excluded by removing cells with less than 500 features and with more than 10% mitochondria gene content. Post alignment, the Seurat objects were merged, normalized with LogNormalize method, and scaled. Top 3000 variable genes were used to anchor. After PCA dimensional reduction with 30 principal components, we mapped our data on UMAP projection with Louvain clustering algorithm. Subsequently canonical marker genes for AT1 and AT2 cells^92^ and sample conditions were used to assign cellular groups.

For RNA velocity, velocyto was used with default run command and run10x sub-command to obtain velocity read counts^93^. For velocity analysis, we used Scanpy and scVelo packages. The datasets were filtered by expression of < 200 genes or > 3200 genes, and genes were expressed in < 3 cells. The dataset was subsequently normalized and the top 3000 highly variable genes with a minimum 20 expressed count threshold were used in preprocessing. A dynamical model was used to recover velocity estimate following Scvelo default parameters (pp.moments(n_neighbors = 30), tl.recover_dynamics(fit_scaling = False), tl.velocity(mode = ’dynamical’)).

We then produced velocity stream overlay on UMAP projection and latent time. We used kmeans clustering from the SKlearn package to identify the two predominant directions of velocity vectors within the nAT2 cell cluster. We then calculated an AT1/AT2 score in these clusters based on expression of cell markers. We further performed partition-based graph abstraction (PAGA)^94^ based on the velocity vectors of each cell type to determine broader cell type linkages^95^.

For STITCH analysis (to generate SPRING plots), we ported in formatted datasets and followed steps of logNormalization, and determined highly variable genes as instructed in the Wagner 2018^96^ example in MATLAB 2023a. The visualization was done using ForcedAtlas layout in Gephi v7.2.1.

For analysis of lung development through embryonic, perinatal, and adult timepoints in published scRNAseq datasets, the processed mRNA counts for each cell were used from developing mouse lung datasets, (GSE119228)^33^, (GSE113320)^12^, and (GSE149563)^37^. Analysis of bleomycin-induced lung injury on AT2 cells was performed on a published scRNAseq dataset (GSE141259)^20^.

### Lung isolation and processing

For prenatal time points, individual embryos were staged by fetal crown–rump length before sacrifice and lungs removed *en bloc*. For postnatal time points, mice were euthanized by either carbon dioxide inhalation or intraperitoneal injection of Pentobarbital (Fatal-Plus Solution) and dissected to both expose the lungs and allow for exsanguination via the abdominal aorta. 5-10 mL of phosphate buffered saline (PBS; Ca^2+^ and Mg^2+^ free, pH 7.4) was gently perfused into the right ventricle of the heart by syringe with a 25-gauge needle until the lungs appeared white. For postnatal time points (≥ 8 days), the trachea was then cannulated with either a blunt 22- or 25- guage catheter, and the lungs were gently inflated to full capacity with molten low melting point agarose (Sigma, 2% in PBS). For all time points, lungs were then separated by lobe and fixed in 4% PFA at 4 °C for either 1 h (e15.5 – e16.5), 1.5 h (e17.5), 2 h (e18.5 – PN10), or 4 h (> PN10). For sectioning, a vibrating microtome (Leica) was used to generate embryonic (150 µm) or adult (250 µm) tissue sections of uniform thickness.

### Immunostaining

Immunohistochemistry was performed as previously described in Treutlein et. al.^9^ using primary antibodies against the following epitopes (used at 1:500 dilution unless otherwise noted): proSPC (rabbit, Sigma AB3786), RAGE (rat, R&D MAB1179), E-cadherin (rat, Life Technologies ECCD-2), PDPN (hamster, DSHB 8.1.1), MUC1 ((hamster, Thermo Scientific HM1630 and rabbit (GeneTex GTX100459)), Alpha-SMA (1:100, mouse IgG2a, Invitrogen 50-9760-82), EZH2 (ENX-1; mouse IgG2b, Santa Cruz Biotechnology sc-166609), KI67 (rat, Thermo Scientific 14-5698-82), RELMα (rabbit, PeproTech 500-P214), CD74 (rat, BD Biosciences 555317), DLK1 (rabbit, R&D Systems MAB8634), C/EBPα (rabbit, Cell Signaling 8178), LAMP1 (rat, DSHB 1D4B-s), and HOPX (rabbit, Abcam ab307671). Briefly, fixed lung slices were permeabilized and blocked overnight in blocking buffer (5% goat serum/PBS/0.5% Triton X-100) at 4 °C. Primary antibody incubation steps were conducted in blocking buffer over 3 nights and washing steps were performed in five rounds (30, 30 60, 90, and 90 min respectively) using the blocking buffer. Primary antibodies were subsequently detected using Alexa Fluor-conjugated secondary antibodies used at 1:1000 dilution unless otherwise noted, including Goat anti-rabbit IgG Alexa 555 conjugated (Invitrogen, A21428), Goat anti-rabbit IgG Alexa 647 conjugated (Invitrogen, A32733), Goat anti-rat IgG Alexa 488 conjugated (Invitrogen, A11006), Goat anti-rat IgG Alexa 647 conjugated (Invitrogen, A21247), Goat anti-hamster Alexa 647 (Invitrogen, A21451), and Goat anti-hamster Alexa 594 (Invitrogen, A21113) unless noted otherwise. Secondary antibody incubation steps were conducted in blocking buffer over 2 nights and washing steps were performed in five rounds (30, 30 60, 90, and 90 min respectively) using PBT (0.1% Tween-20/PBS). For co-staining using antibodies from the same rabbit origins, we performed sequential staining with 1-day anti-rabbit blocking between processes. In brief, we stain with primary and secondary antibodies of the 1^st^ marker as described, and then we block the stained tissue with 40 μg/mL Fab Fragment Goat Anti-Rabbit IgG (Jackson ImmunoResearch, AffiniPure™, 111-007-008) prior to the 2^nd^ marker staining. EdU was detected using the standard Click-iT reaction (Thermo Fisher). Lung slices were then mounted in Vectashield Mounting Medium (Vector labs, H1000) and sealed with a coverslip. Images were acquired using a laser-scanning confocal microscope (Zeiss LSM 780 and Leica Stellaris 8) and subsequently processed using ImageJ.

### Timelapse Imaging

Fluorescently lineage labeled mouse lung were sectioned at 200 μm thick using compresstome. When imaging, we image sectioned slices in serum-free DMEM media on top of Matrigel for stability, using Timelapse setup in LAS-X software for 24 h consecutively.

### Confocal Images 3D reconstruction

We reconstructed 3D representation of cell imaging using IMARIS 10.2.1. The z-stack images are imported as .tif images and reconstructed into 3D representation of tissue images. Marker intensities are individually normalized thresholds for intensity and contrast, smoothing cellular structures for interpretation accuracy, and removal background noise.

### Cell isolation and culture

For time points e15.5, e18.5, and PN3, lungs were isolated *en bloc* without perfusion and pooled by litter (6-8 lungs) for further processing. The lungs were then dissociated in 50 U/mL Dispase (Corning 354235) and triturated with flame sterilized glass Pasteur pipettes of decreasing diameters (1,000 µm, 800 µm, and 600 µm) to achieve a single-cell suspension. For PN25 and adult time points, the lung was perfused with DPBS (Ca^2+^ and Mg^2+^ free, pH 7.4) as described above. The trachea was punctured and then cannulated with a blunt 22-guage catheter, and the lungs were inflated to full capacity with Dispase (∼1 mL), and the trachea was sealed using a non-absorbable silk suture. The inflated lobes were incubated in Dispase for 45 min at 25 °C on a shaker.

For all time points, digested lungs were minced with a razor blade, suspended in 7 mL of digestion buffer (RPMI-1640 containing FBS (10%, Corning), penicillin-streptomycin (1 U/mL, Thermo Scientific), and DNase I (0.33 U/mL, Roche)), incubated at 4 °C for 10 min on a flat shaker and triturated briefly with a 5-mL pipette.

To deplete red blood cells, an equal volume of AT2 isolation media (RPMI-1640 supplemented with penicillin-streptomycin (1 U/mL, Thermo Scientific), and DNase I (0.33 U/mL, Roche)) was added to the lung single-cell suspension. The suspension was passed through a 100 µm mesh filter to remove residual tissue fragments and centrifuged at 400×g, 4 °C for 10 min. Pelleted cells were resuspended in 5 mL of red blood lysis buffer (BD Biosciences), incubated for 2 min at room temperature, and neutralized with an equal volume of AT2 isolation media. The suspension was passed through a 40 µm mesh filter (Fisher), centrifuged at 400×g, 4 °C for 10 min, and then resuspended in MACS buffer (2 mM EDTA, 0.5% BSA in PBS, filtered and degassed) for purification.

The resultant single-cell suspension was incubated with Mouse Fc Blocking Reagent (Miltenyi Biotec, 130-092-575) for 5 min at 4 °C. Other cell types were first depleted with antibodies against CD45 (eBioscience, 13-0451-85), CD31 (BD Biosciences, 557355), Ter119 (eBioscience, 13-5921-85), CD140a (Miltenyi Biotec, 130-101-905), and CD104 (Miltenyi Biotec, 130-106-922), followed by anti-biotin (Miltenyi Biotec, 130-090-485) magnetic MicroBeads. Cell suspensions were passed through a 35 µm cell strainer (BD Biosciences) prior to being loaded on LD columns (Miltenyi Biotec) as per the vendor’s protocol. The flow through was centrifuged at 300×g, 4 °C for 5 min and the pelleted cells were resuspended in MACS buffer.

For positive selection of cell types, the following antibodies were used: anti-EpCAM (clone G8.8, eBioscience, 13-5791-82) for e15.5 distal bipotent progenitors, anti-RELMα (PeproTech, 500-P214BT) for e18.5 nascent AT2 cells, and either anti-CD74 or anti-MHC-II (clone M5/114.15.2, eBioscience, 13-5321-82) for mature AT2 cells. Following antibody incubation, the cells were then incubated with either anti-biotin or anti-rat IgG MicroBeads and loaded on MS columns (Miltenyi Biotec) as per the vendor’s protocol. The flow-through was discarded and the column was taken off the magnet. The cells remaining in the column were eluted with MACS buffer, resulting in a highly enriched preparation of either bipotent progenitors (EpCAM^+^), nascent AT2s (RELMɑ^+^), or mature AT2s (MHC-II^+^), unless noted otherwise.

For culturing bipotent progenitors, nAT2s, or mAT2s, cell density and viability were calculated using a Vi-CELL XR Cell Viability Analyzer (Beckman Coulter). Cells were cultured in eight-well #1.5 coverglass chambers (Cellvis, C8-1.5H-N), pre-coated with growth factor reduced Matrigel (85 µL, Corning 354230) for 30 min at 37 °C. The cells were plated at varying densities: 30,000 cells/well (for e15.5 and e18.5), 60,000 cells/well for PN3), and 80,000 cells/well (for PN25 and adult). Cells were supplemented with FGF7 (50 ng/mL, R&D Systems 5028-KG-025), GSK126 (10 µM, ApexBio A3446), KL001 (12 µM, MedChemExpress HY-108468), and interferon-γ (20 ng/mL, PeproTech 315-05) to the indicated concentrations in 400 µL of DMEM/F12 supplemented with penicillin-streptomycin (1 U/mL, Thermo Scientific).

Excluding FGF7, all other small molecules listed above were added after a 12h incubation period to allow for cells to seed and adhere to the layer of Matrigel. Cells were maintained in an incubator at 37 °C with 5% CO2/air, with media changes every other day typically for 4 days, except where indicated otherwise.

### Flow cytometry

We perform flow cytometry of AT2 cells using Miltenyl’s Tyto flow sorter. Filtered and RBC depleted lung cell isolates are to be incubated with 20 µL of each of the following 1:50 diluted primary antibodies: anti-RELMɑ, anti-CD74, and anti-MHC-II for 30 min at 4 °C. Cells are washed with sorting buffer (Miltenyi, 130-107-207) and subsequently incubated with 20 µL of each of the following secondary antibodies: (Anti-rabbit Alexa Fluor 647, anti-rat Alexa Fluor 405) for 30 min at 4 °C. Subsequently the sample are washed, filtered, and resuspended in 1 mL sorting buffer and transfer to the sorting compartment of the cartridge (Miltenyi, 130-104-791). Gates for markers positivity are drawn using unstained cells as negative controls.

### BH3 analysis

Lung samples from mice of different ages (e15.5, e18.5, and ≥ PN60) were dissociated into a single-cell suspension and enriched using relevant markers as described above. Cells were stained on ice for 25 min away from light, then centrifuged at 200×g for 5 min, and subjected to flow cytometry–based BH3 profiling as previously described by the Sarosiek lab^78^. Briefly, cells were treated with activator or sensitizer BH3 peptides (New England Peptide) for 60 min at 28 °C in mannitol experimental buffer (MEB) [10 mM Hepes (pH 7.5), 150 mM mannitol, 50 mM KCl, 0.02 mM EGTA, 0.02 mM EDTA, 0.1% BSA, and 5 mM succinate] with 0.001% digitonin. Peptide sequences are as follows: BIM (Ac-MRPEIWIAQELRRIGDEFNA-NH2), BID (Ac-EDIIRNIARHLAQVGDSMDRY-NH2), PUMA (p53 upregulated modulator of apoptosis) (Ac-EQWAREIGAQLRRMADDLNA-NH2), BAD (Ac-LWAAQRYGRELRRMSDEFEGSFKGL-NH2), Hrk (Harakiri) (Ac-WSSAAQLTAARLKALGDELHQ-NH2), and MS1 (MCL-1 specific) (Ac-RPEIWMTQGLRRLGDEINAYYAR-NH2). After peptide exposure, cells were fixed in 2% PFA for 15 min which was then neutralized by addition of N2 buffer [1.7 M tris base and 1.25 M glycine (pH 9.1)]. Cells were stained overnight with DAPI (1:1000, Abcam) and anti–cytochrome c–Alexa Fluor 647 (1:2000, clone 6H2.B4, BioLegend) in a saponin-based buffer (final concentration, 0.1% saponin; 1% BSA) and then analyzed by flow cytometry.

Cytochrome c release in response to BIM treatment was measured on an Attune NxT flow cytometer (Thermo Fisher Scientific).

### Conditional deletion of *Cebpa* and *Ezh2 in vivo*

For conditional and efficient recombination of floxed alleles (*Cebpa^flox^*or *Ezh2^flox^*) at different time points and cell types, we bred the floxed allele into one of four mouse lines, each with a distinct combination of Cre and reporter alleles. For in vivo sparse cell labeling with efficient recombination of either *Cebpa^flox^* or *Ezh2^flo^*alleles in distal progenitors, the *Tg^Nkx^*^2^.^1^*^-Cre^*; *Rosa26^mTmG/mTmG^* mouse line was used as we have previously found it sparsely labels DPs with efficient flox allele recombination^10^. For sparse labeling of nAT2s with efficient recombination of *Cebpa^flox^*, the *LyzM^Cre^*; *Rosa26^mTmG/mTmG^* mouse line was used as previously described^10^. For broad labeling of AT2 cells with efficient *Cebpa^flox^* recombination following a high dose of tamoxifen (20 mg/kg), the *Sftpc^Cre-ERT^*^2^; *Rosa26^mTmG/mTmG^* mouse line was used. Finally, to sparsely label AT2s and retain efficient *Cebpa^flox^* recombination following a low dose of tamoxifen (30 ng/kg), the *Sftpc^Cre-ERT^*; *Tg(iSuRe-Cre)* mouse line was used. Upon its recombination, the *Tg(iSuRe-Cre)* allele constitutively expresses both the fluorescent lineage label tdTomato as well as Cre recombinase, an approach that has been well established to efficiently recombine multiple floxed under lower tamoxifen doses for targeting sparse cells^49^. To determine proliferation of AT2 cells, EdU (1 mg/mL) was administered in the drinking water for the indicated period of time.

### ATAC-seq and CHIP-seq analysis

Cells from PN25 *Sftpc^CreER^; Tg(iSuRe-Cre); Cebpa^wt/fl^* and *Sftpc^CreER^; Tg(iSuRe-Cre); Cebpa^fl/fl^* mice (2 weeks after TMX) were isolated and enriched (EpCAM^+^) as per protocol described above. Raw sequencing reads were trimmed of sequencing adapters and bad quality bases with cutadapt version 2.8 in Trim Galore version 0.6.6 using default parameters. Bowtie2 version 2.5.0 (parameters: --end-to-end --very-sensitive --no-mixed --no-discordant --phred33 -I 10 -X 700) was used to map trimmed reads to the reference genome Mus_musculus.GRCm38. Picard MarkDuplicates (v.2.26.10; http://broadinstitute.github.io/picard) was used for duplicate removal. Reads were shifted by + 4 bp for those mapping to the positive strand and ° 5 bp for those mapping to the negative strand. ATAC-seq analysis was performed with HOMER(v 4.11). Tag directories were made for each ATAC-seq run using HOMER^97^ “makeTagDirectory”. Peaks for individual replicates were identified using the “findPeaks” command. To generate high-confidence peak lists across both replicate samples for downstream analysis, ATAC-seq peaks were identified using the HOMER “getDifferentialPeaksReplicates.pl” command. This command uses DESeq2^98, 99^ and identifies peaks that pass 2-fold enrichment and false discovery rate (FDR) < 0.05 cutoffs. For these analyses, peak lists combining all peaks from the comparison conditions were assembled. One condition was used as the target, and the other was used as background. To characterize enriched motifs near differentially enriched peaks, we used HOMER’s “findMotifsGenome.pl” command to scan the entire peak by using “-size given” with standard background. ATAC-seq peak annotation was performed using the homer annotatePeaks.pl function to assign the nearest gene name to the peaks. To estimate the enrichment of transcription factors and epigenetic modifications on the murine genome, we used the Cistrome DB Toolkit analysis browser^61, 100^ using default settings.

### Incoherent feed forward loop type 2 model fitting

To assess whether the temporal expression pattern of *Dlk1* fit that predicted by a feed forward loop we used the publicly available biocircuits package (https://biocircuits.github.io/) to numerically solve the differential equations for a type 2, incoherent feed forward loop following a step reduction of PRC2. After solving for a function over time for *Dlk1*, an R script was generated to align its peak value to that observed in the scRNAseq timecourse and calculate a goodness of fit by root mean squared error (RMSE). Parameters of the I2-FFL model were independently varied until a good fit was reached (beta = 1.9, gamma = 5.5, and kappa = 3).

### Mouse lung injury experiments and viral transfections

Lung injury was induced in adult mice by single-dose administration of bleomycin sulfate (0.25mg/kg; VWR J60727-MA), which was dissolved in sterile saline and administered via intratracheal (i.t.) administration. Lungs were isolated 16 days post injury and processed for immunostaining.

To test whether C/EBPα is required for proper injury repair, adult *Rosa26^mTmG/mTmG^* mice were administered either an AAV-*Sftpc-Cre* (control), or an AAV-*Sftpc-Cebpa-T2A-Cre* (prepared by the Boston Children’s Hospital Viral Vector Core (1.5 × 10^14^ GC (genome copies)/mL, serotype 2/9) and intratracheally instilled (5 µL of viral solution diluted in 45 µL PBS)) to overdrive *Cebpa* expression and induce GFP labeling of AT2 lineage. A minimal 320 bp *Sftpc* promoter element was used for both AAV preps. 5 days later, all mice were administered Fiin1 hydrochloride (14.75 mg/kg; Tocris 4002) via intratracheal instillation as described in out prior work^10^ or BHT (225 mg/kg; Sigma B1378) via intraperitoneal injection^71^ to induce lung injury. Lungs were isolated 14 days post injury and processed for immunostaining. For all intratracheal instillation experiments, mice were anesthetized with isoflurane and kept in recovery under a heat lamp.

### Small molecule induction of CHOP

To induce CHOP expression in AT2 cells, *Sftpc^CreERT^*^2^*^-rtTA^; Rosa26^mTmG/mTmG^* mice were first administered tamoxifen (20 mg/kg) at PN11 for labeling of AT2 cells with GFP. 24 h later, mice were either given corn oil (vehicle) or CCT020312 (MedChem Express HY-119240), a PERK activator found to upregulate CHOP expression^101^, at 2 mg/kg dissolved in corn oil via intraperitoneal injection every 4 days (4 doses total). Lungs were isolated 2 weeks later and processed for immunostaining.

### Statistical analysis

Data analysis and all statistical tests were performed with GraphPad Prism software or R. Replicate experiments were biological replicated using different animals, and the quantitative values are presented as mean ± SD unless indicated otherwise. In graphs displayed as box plots, the horizontal lines represent the median and error bars depict the 75^th^ – 25^th^ interquartile range (IQR). Two-sided Student’s *t*-tests (for normally distributed data), Mann-Whitney U-tests (for data not normally distributed), and Brown-Forsythe and Welch ANOVA (for comparisons between ≥ 3 groups) with Dunnett’s multiple comparisons test were used to determine P-values. No statistical method was used to predetermine sample size, and data distribution was tested for normality prior to statistical analysis and plotting.

## Data Availability

The authors confirm that all relevant data are available in the paper or its Supplementary Information files. The scRNAseq datasets are available in the GEO repository under accession numbers “GSE119228”, “GSE113320”, “GSE109774”, and “GSE149563” for developing and adult mouse lung cells, “GSE295269” for CEBP/α knockout mouse lung epithelial cells, “GSE141259” for adult bleomycin injury, “GSE218666” for AT1 injury, “GSE184384” for influenza injury, and “GSE113049” for LPS injury models. Source Data are available for this paper.

### Code Availability

Code to reproduce the analyses described in this manuscript can be accessed at https://github.com/JimmyMayoGit/Sawhney_2025

## Acknowledgments

The authors thank members of the Brownfield laboratory for helpful discussions and critical reading of the manuscript. We thank D. Tenen (*Cebpa^fl/fl^*) for the transgenic mouse strain; Hannah Golding for performing organotypic culture experiments; and Gamaliel Taengwa for data quantification. This work was supported by R00HL127267, R01HL171056, and a Pilot Award from Mayo Clinic’s Center for Biomedical Discovery (D.G.B).

## Author Contributions

ASS, BJD, and DGB conceived the study and designed the experiments. ASS, BJD, JC, DG, GA, AO, HH, LS, XQ, SRH, ASU performed the experiments. ASS, BJD, and AO analyzed and interpreted the data. JC, AH, EL, and DGB performed the scRNAseq analysis. DG and ZW performed the ATACseq analysis. RW and SD performed bleomycin injury experiments, and KAS and ZI performed BH3 profiling experiments. DGB, ASS, and BJD wrote the paper.

## Ethics Declarations

### Competing interests

The authors have no competing interests to declare.

---

**Supplementary Figure 1:**
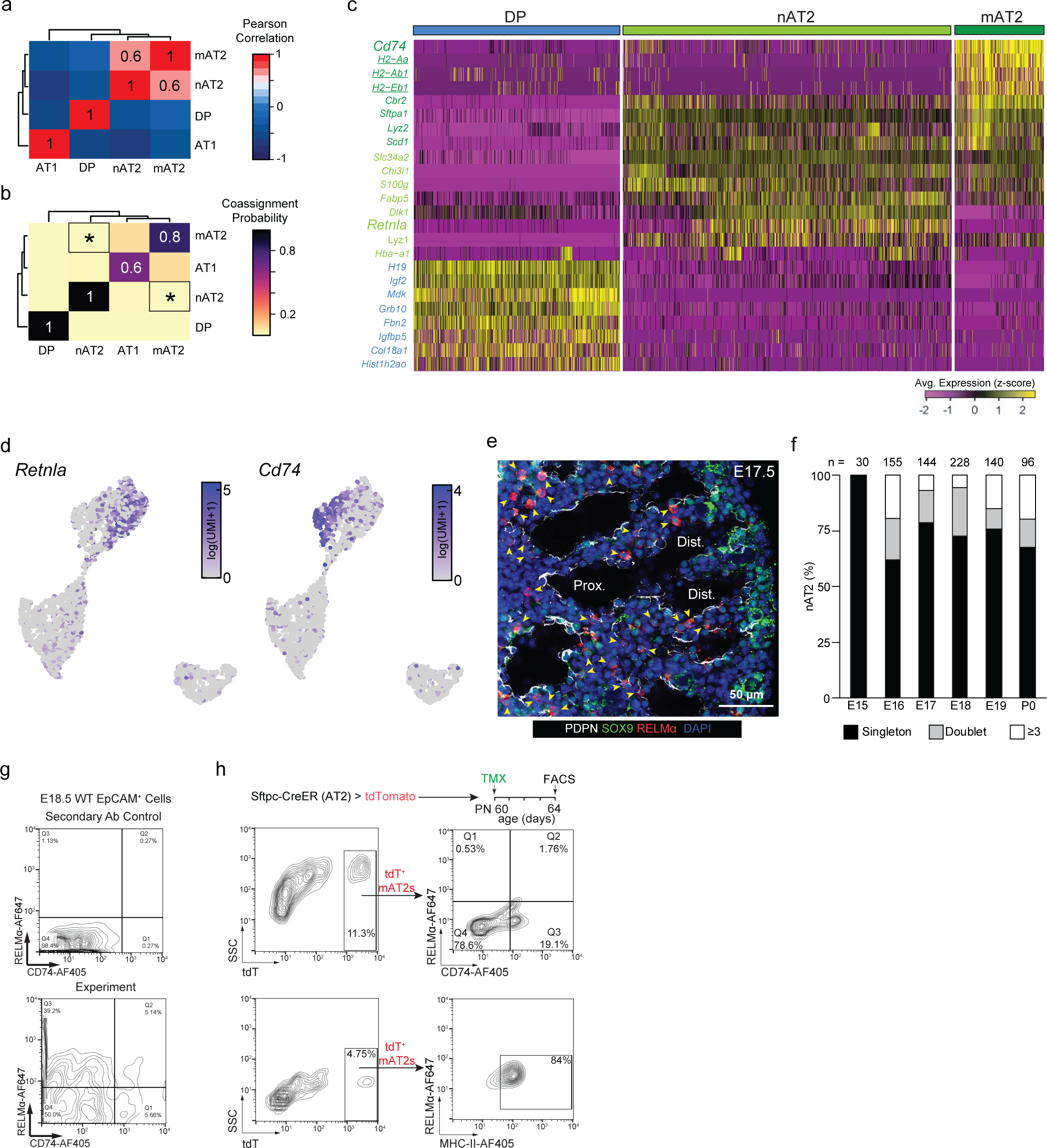
Dynamics of nascent AT2 state and emergence. **a** Cluster correlation analyzing the strength of correlation between each of the four clusters detected: AT1, DP, nAT2, and mAT2. Pearson Correlation > 0.5 indicates a strong, positive correlation. **b** Coassignment probability for the four clusters shown in (**a**), measuring the probability of merging two clusters. Note that the nAT2 and mAT2 clusters do not have a significant coassignment probability (asterisk), indicating they are distinct, stable cell populations. Low coassignment probability (< 0.5) represents a stable cluster with respect to one another. **c** scRNAseq expression for genes enriched for the clusters identified within the AT2 lineage. *Cd74* and *Retnla* are highlighted (bold) as being surface markers chosen for mature and nascent AT2s respectively. **d** feature plot of *Retnla* and *Cd74* in mouse development timecourse dataset from Cohen et al., showing high and exclusive expression patterns of the nascent and mature AT2 cell markers. **e** e17.5 lung immunostained for PDPN (white), distal progenitor marker SOX9 (green), nAT2 marker RELMα (red), and DAPI (blue). Nascent AT2 cells are marked (arrowhead) to show them as RELMα^+^ and SOX9^low^. **f** Quantification of nAT2 clustering observed as either singlets, doublets, or in clusters of ≥ 3 across embryonic timepoints from e15.5 to PN0 (*n* = 30 nAT2 clusters scored at e15.5, 155 at e16.5, 144 at e17.5, 228 at e18.5, 140 at e19.5, and 96 at P0). **g** flow cytometry of EpCAM^+^ nascent AT2s from *WT embryonic* mouse lung, show moderate RELMα positivity, and low labeling efficiency by CD74, suggesting that nascent AT2s can be labeled and sort by RELMα. **h** flow cytometry of tdT^+^ mature AT2s from *Sftpc^CreER^; Tg(iSuRe-Cre)*mouse lung, show little RELMα positivity, moderate labeling efficiency by CD74, and high labeling efficiency by MHC-II, suggesting that MHC-II is more effective as cell isolating marker comparing to CD74.

**Supplementary Figure 2:**
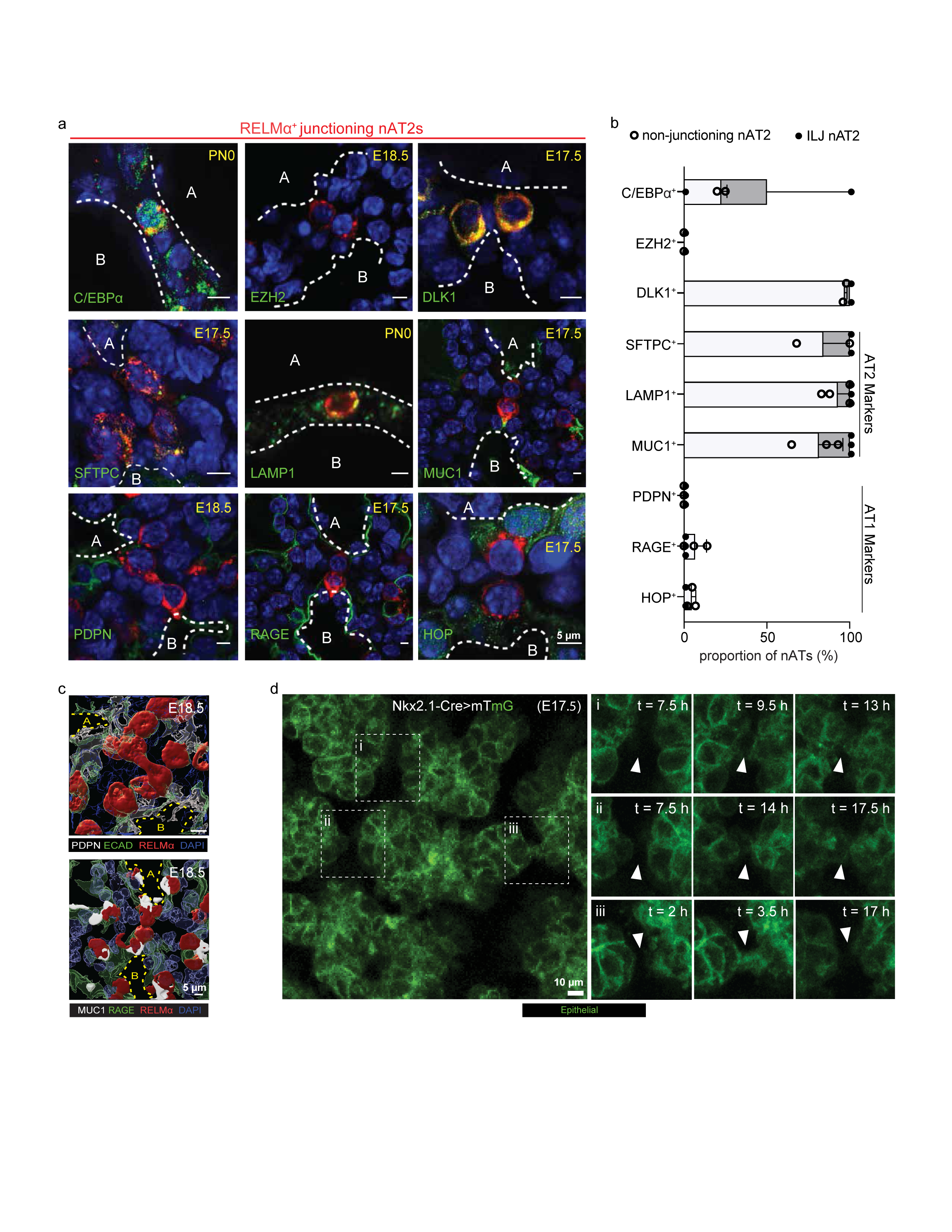
Spatial and Temporal Characterization of junctioning nAT2s. **a** Cross-section of e17.5, e18.5, or PN0 distal lungs slices immunostained for RELMα (red), DAPI (blue), and markers of interests (green), including C/EBPα, EZH2, DLK1, SFTPC, LAMP1, MUC1, PDPN, RAGE, and HOP. **b** Quantification of portion of nAT2 showing their high positivity in AT2 character and fate plasticity and their absence of AT1 character. **c** Additional 3D renderings on immunostaining of double interlumenal junctioning nAT2s for E-cadherin (ECAD) or RAGE (green), RELMα (red), PDPN or MUC1 (white), and DAPI (blue). Lumen boundaries are marked in yellow dashes. **d** Additional snapshots timeplase confocal microscopy on *Nkx2.1^Cre^*; *Rosa26^mTmG^* lineage labeled precision cut lung slices at distal lung showing dynamics interlumenal junctioning.

**Supplementary Figure 3:**
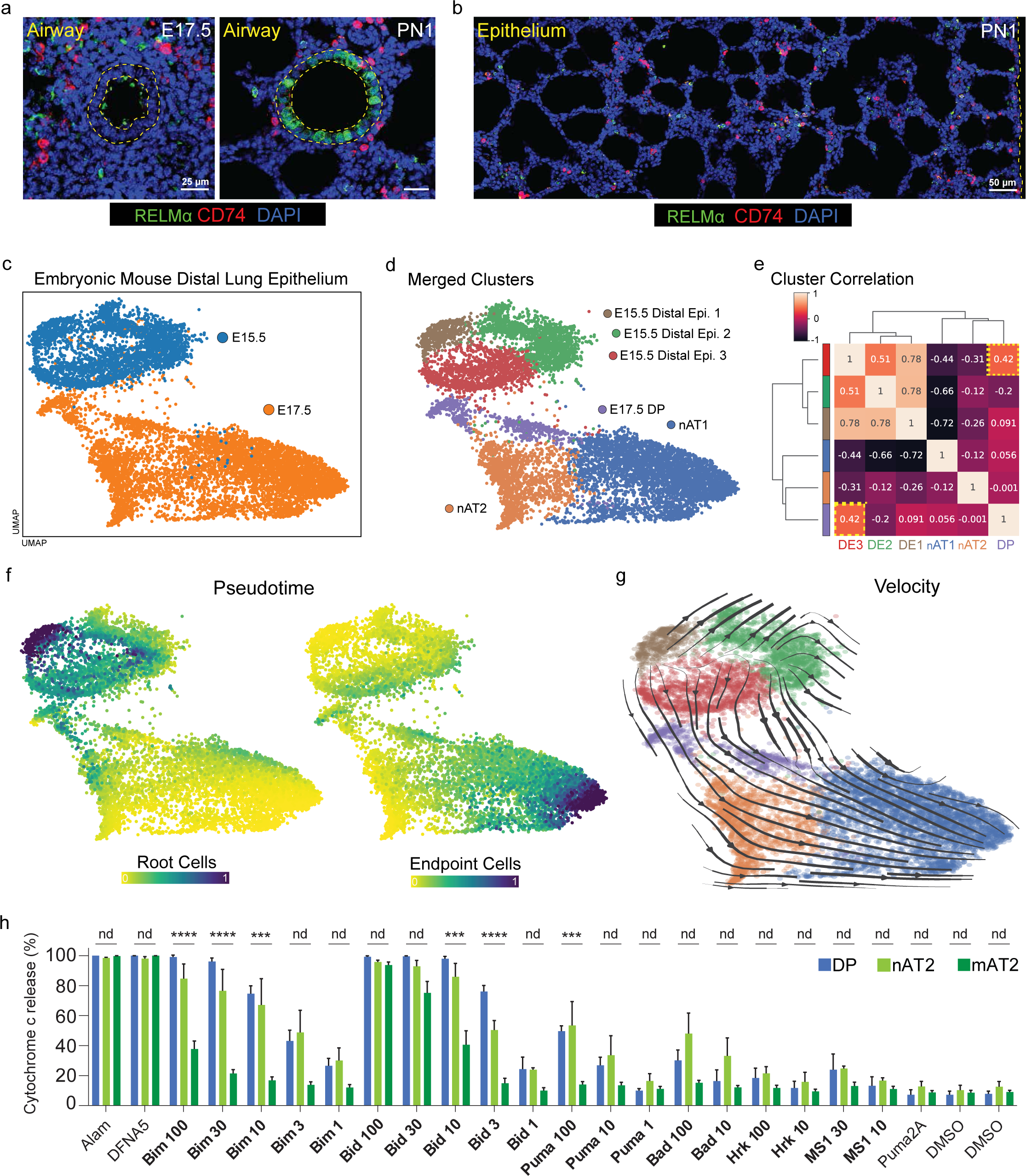
Details on state dependent plasticity and death sensitivity. **a** Cross-section of an airway (dashed yellow) of e17.5 (left) and PN1 (right) lungs immunostained for RELMα (green), CD74 (red) and DAPI (blue). Airway region at PN1 shows presence of RELMα in the airway epithelium, but the absence at e17.5 confirms its specificity to nAT2s during the AT2 fate selection window. Bars, 25 µm. **b** Proximal and distal epithelium of a PN1 lung immunostained for RELMα (green), CD74 (red) and DAPI (blue), showing salt-and-pepper pattern of AT2 maturation. Bar, 50 µm. **c** scRNAseq UMAP of distal lung epithelium at e15.5 (blue) and e17.5 (orange) timepoints. **d** Merged clusters of scRNAseq dataset in (**c**), showing three e15.5 distal epithelium (DE) clusters, and three e17.5 clusters (DP, nAT2, and nAT1). **e** Cluster correlation of the six clusters observed in (**d**). No significant correlation observed between either e15.5 DE cluster with the nAT2 or nAT1 clusters at e17.5. A weak correlation (0.42, dashed box) observed between the e15.5 DE3 and e17.5 DP clusters. **f** Pseudotime trajectory analysis showing e15.5 cells are the primary root cells of origin (left) and e17.5 AT1 cells are endpoint cells (right). **g** Velocity trajectories showing transition from e15.5 DE clusters to e17.5 DP cells. Subsequent trajectories lead either directly to AT1 cells or indirectly, transitioning through the nAT2 cluster. **h** BH3 death sensitivity assay of e15.5 DP, e18.5 nAT2, and ≥ PN60 mAT2 cells, showing percent cytochrome c release upon treatment with activator or sensitizer BH3 peptides. A higher cytochrome c release percentage indicates increased sensitivity to BH3-mediated apoptosis. Alam and DFNA5 are positive controls; DMSO is the vehicle control and PUMA 2A is an inactive peptide negative control. Cytochrome c release (%) values are presented as mean ± SD. \*\*\*\**p* ≤ 0.0001, \*\*\**p* ≤ 0.001 (one-way ANOVA, comparing DP and mAT2 cells).

**Supplementary Figure 4:**
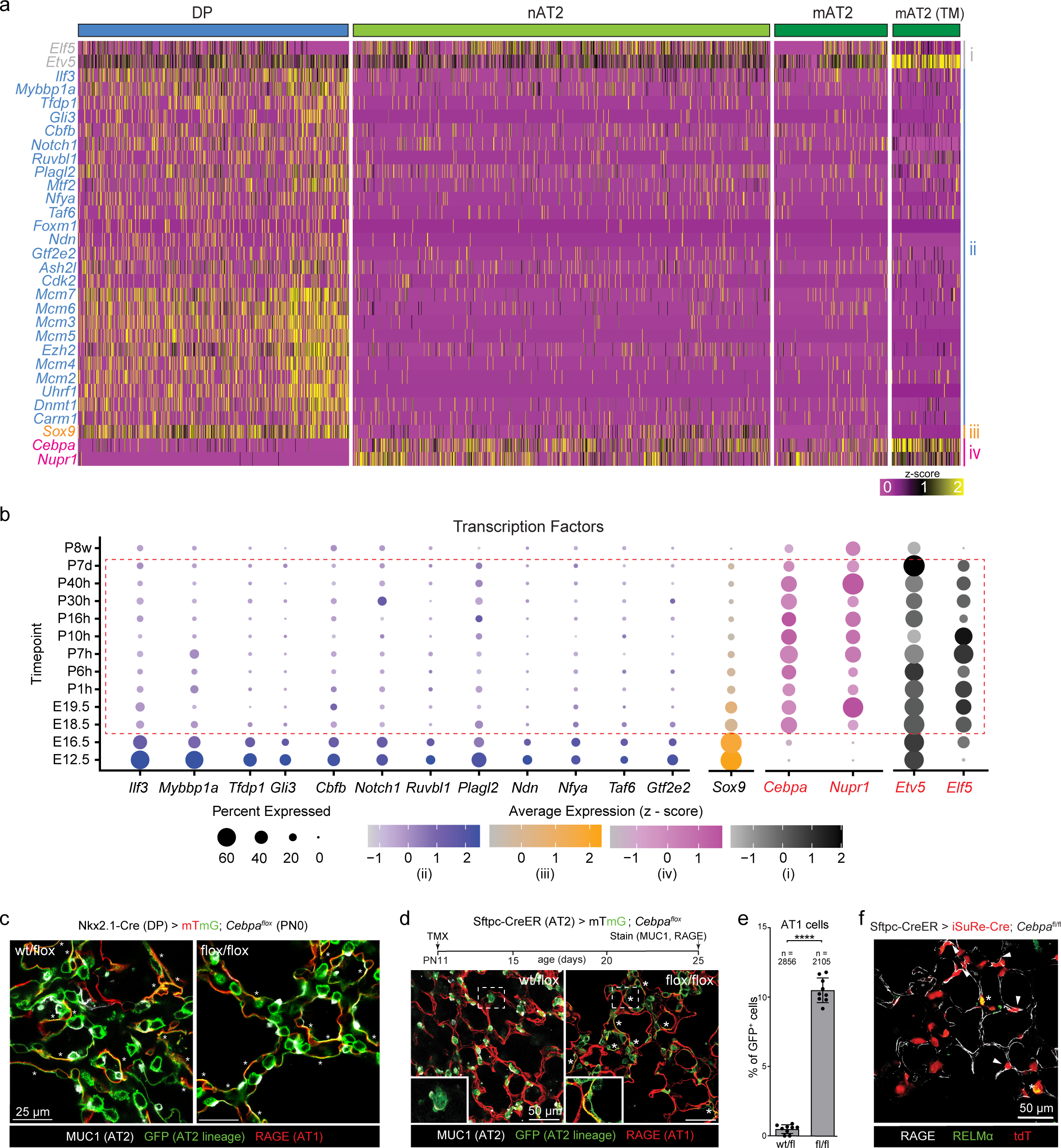
Bioinformatic screen for regulators of AT2 plasticity and perinatal *Cebpa* deletion. **a** Heat map showing expression of all 32 genes from Figure 4a. Cells (columns) are arranged in developmental pseudotime (DP, nAT2, mAT2; left to right) along the AT2 lineage. Note that the “mAT2 (TM)” is from the Tabula Muris scRNAseq dataset (GSE109774)^41^, which sequences AT2 cells more deeply. Genes form four patterns: constitutive (grey), DP restrictive (blue), DP-nAT2 restrictive (orange), and nAT2-mAT2 restrictive (pink). **b** Dot plot of the transcription factors from (**a**) that are expressed within the plasticity window (red dotted box), showing expression at different developmental and adult timepoints. **c** Alveolar regions of PN0 control *Nkx2.1^Cre^; Rosa26^mTmG^; Cebpa^wt/fl^*(left) and *Nkx2.1^Cre^; Rosa26^mTmG^; Cebpa^fl/fl^*(right) lungs immunostained for MUC1 (white) and RAGE (red). GFP^+^ (DP lineage) cells form both AT1 cells (flattening; asterisk “*”) and AT2 cells (MUC1^+^) even upon *Cebpa* deletion. Bars, 25 µm. **d** Experimental design (top) and immunostains (bottom) for simultaneous *Cebpa* deletion and lineage labeling AT2 cells at PN11 via intraperitoneal (i.p.) tamoxifen (TMX) induction. Lungs were isolated 14 days post TMX (PN25) and immunostained for MUC1 (white) and RAGE (red). Alveolar regions of control *Sftpc^CreER^; Rosa26^mTmG^; Cebpa^wt/fl^* (left) and *Sftpc^CreER^; Rosa26^mTmG^; Cebpa^fl/fl^* (right) lungs immunostained as indicated, showing GFP^+^/RAGE^+^ AT1 cells (asterisk) upon *Cebpa* deletion. Close up (bottom left) panels for each condition show a GFP^+^/MUC1^+^ AT2 (control; left) or GFP^+^/RAGE^+^ AT1 cell (*Cebpa* deletion; right). Bars, 100 µm. **e** Quantification of (**d**) showing percent GFP^+^ cells that differentiated into AT1s upon *Cebpa* deletion (*n* = 2856 GFP^+^ cells sampled for control and 2105 for *Cebpa* deletion in experimental triplicate). \*\*\*\**p* = 6.4 × 10^-16^ (Student’s two-sided *t*-test, data as mean ± SD). All experiments were repeated at least three times. **f** Alveolar region of *Sftpc^CreER^; Tg(iSuRe-Cre); Cebpa^fl/fl^* mouse lung immunostained for RAGE (white), RELMα (green) and Cre reporter (tdT). Lineage labeled RAGE^+^ (arrowheads) and RELMα^+^ (asterisk) cells are mutually exclusive from one another, showing the two distinct states upon loss of *Cebpa*.

**Supplementary Figure 5:**
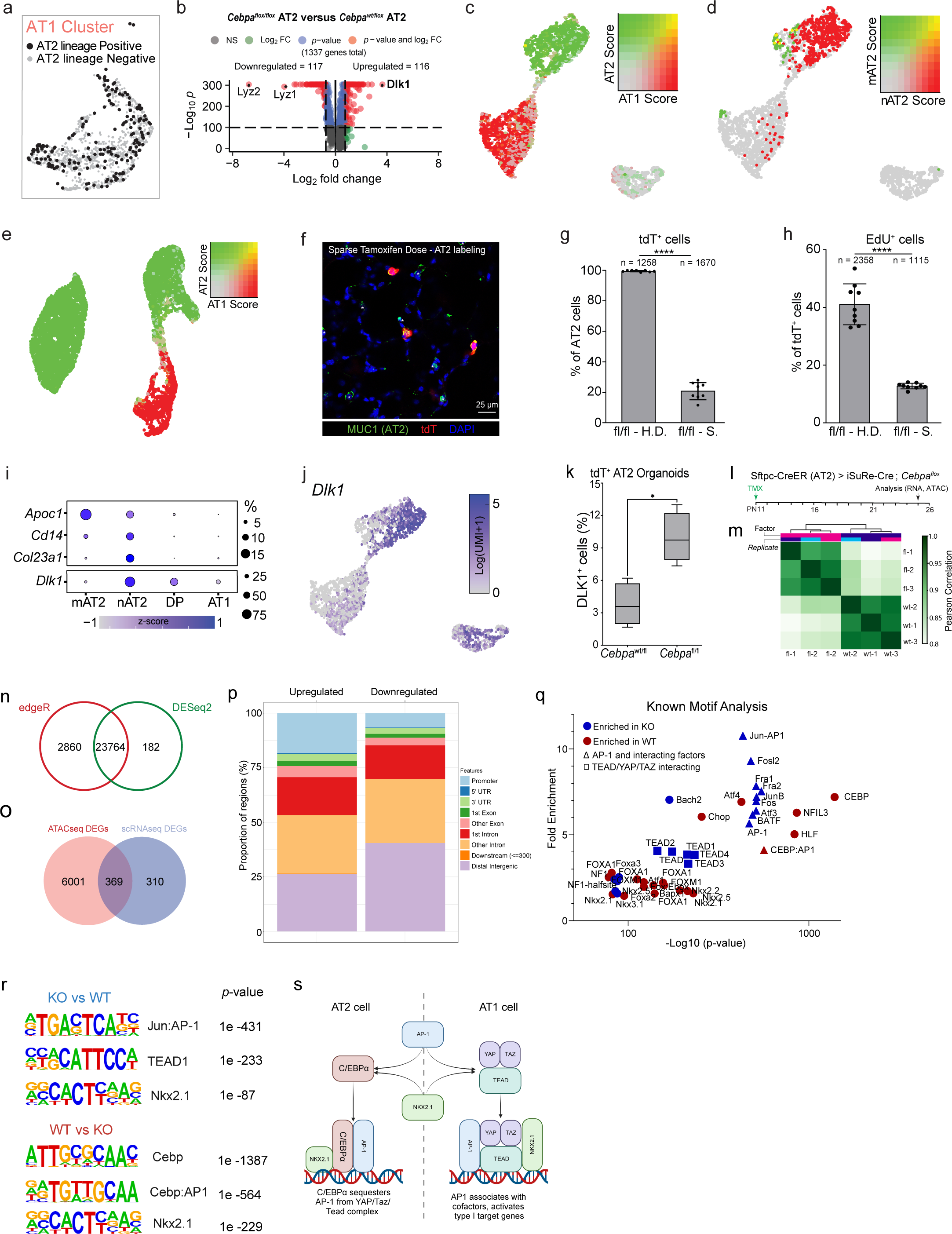
*Cebpa* deletion and gene score validation. **a** scRNAseq UMAP close-up of AT1 cluster (Figure 5a-i) depicting which cells are lineage positive (black) that converted to AT1s upon deletion of *Cebpa*. **b** Volcano plot showing downregulated (left) and upregulated (right) genes in AT2 cells upon *Cebpa* deletion. Genes are considered to have significant down/upregulation if –Log_10_ *P* > 100, and Log_2_ fold change > 1 or < – 1 (red). **c** AT1 and AT2 gene score co-expression on scRNAseq UMAP of the developing lung^33^ to assess changes in either AT2 or AT1 related genes. **d** nAT2 and mAT2 gene score co-expression on scRNAseq UMAP of the developing lung^33^ to confirm restrictive expression of genes in either AT2 state. **e** AT1 and AT2 gene score (see Suppl. Table 1) co-expression on scRNAseq UMAP, showing retained AT2 gene score, and lack of an AT1 score upon loss of *Cebpa* in AT2 cells. **f** Alveolar regions of a PN25 *Sftpc^CreER^; Tg(iSuRe-Cre)*mouse lung, 14 days after Cre induction with a sparse dose of TMX. Lungs were immunostained for MUC1 (green), tdT (red), and DAPI (blue). Bar, 25 µm. **g** Quantification of (**f**) for percent tdT^+^ labeling of AT2 cells in either sparse or high doses of TMX (*n* = 1258 AT2 cells sampled for high dose TMX and 1670 for the sparse dose TMX). \*\*\*\**p* = 1.73 × 10^-9^ (Student’s two-sided *t*-test, data as mean ± SD). **h** Quantification of Figure 5b, showing percent tdT^+^/EdU^+^ proliferating cells upon *Cebpa* deletion in either High Dose or Sparse conditions. Note a significant reduction in proliferation upon sparse *Cebpa* deletion (*n* = 2358 tdT^+^ cells sampled for the high dose deletion and 1115 for the sparse deletion in experimental triplicate). \*\*\*\**p* = 2.47 × 10^-9^ (Student’s two-sided *t*-test, data as mean ± SD). **i** Dot plot showing expression of the six genes (encoding for extracellular proteins) upregulated in the *Cebpa^fl/fl^* cluster, also enriched in nAT2 cells (Figure 5d). **j** *Dlk1* gene expression on a scRNAseq developmental dataset confirming DLk1 expression in mAT2 cells. **k** Quantification of Lineage labeled organoids generated from AT2s isolated from *Sftpc^CreER^; Tg(iSuRe-Cre); Cebpa^wt/fl^* and *Sftpc^CreER^; Tg(iSuRe-Cre); Cebpa^fl/fl^* murine lung, showing higher percent DLK1^+^ AT2s cells upon *Cebpa* deletion (*n* = 435 tdT^+^ cells sampled for the *Cebpa* deletion and 435 for control in experimental triplicate). \*\*\*\**p* = 0.0388 (Student’s two-sided *t*-test). **l** Experimental design for labeling and deleting *Cebpa* in AT2 cells of *Sftpc-CreER;Tg(iSuRe-Cre);Cebpa^wt/fl^* and *Sftpc-CreER;Tg(iSuRe-Cre);Cebpa^fl/fl^*mice at PN11, isolating cells 14 days post TMX (at PN25) for scRNAseq and bulk ATACseq analyses. **m** Hierarchical clustering of sample binding sites shows high correlation between conditions in the ATAC-seq data. **n** Venn diagram of binding site overlaps in the ATAC-seq data between two programs for differential accessibility analysis – edgeR and DESeq2. **o** Differentially expressed gene (DEG) overlaps between RNA-seq and ATAC-seq data for the *Cebpa^fl/fl^* vs *Cebpa^wt/fl^*AT2 cells. **p** Feature distribution from the ATAC-seq data detailing the genomic features associated with down-and upregulated binding sites upon loss of *Cebpa*. **q** HOMER analysis of the ATAC-seq data shows motifs enriched in the *Cebpa^wt/fl^*(red) and *Cebpa^fl/fl^* (blue). AP-1 and Yap/Taz interacting factors are highlighted via shape. **r** HOMER motifs and associated *p*-values highlight a role of AP1, TEAD, and NKX2.1 signaling in both the WT and KO conditions. **s** Schematic detailing the potential role of C/EBPα interacting with AP1 and NKX2.1 in AT2 (left) and AT1 (right) cells.

**Supplementary Figure 6:**
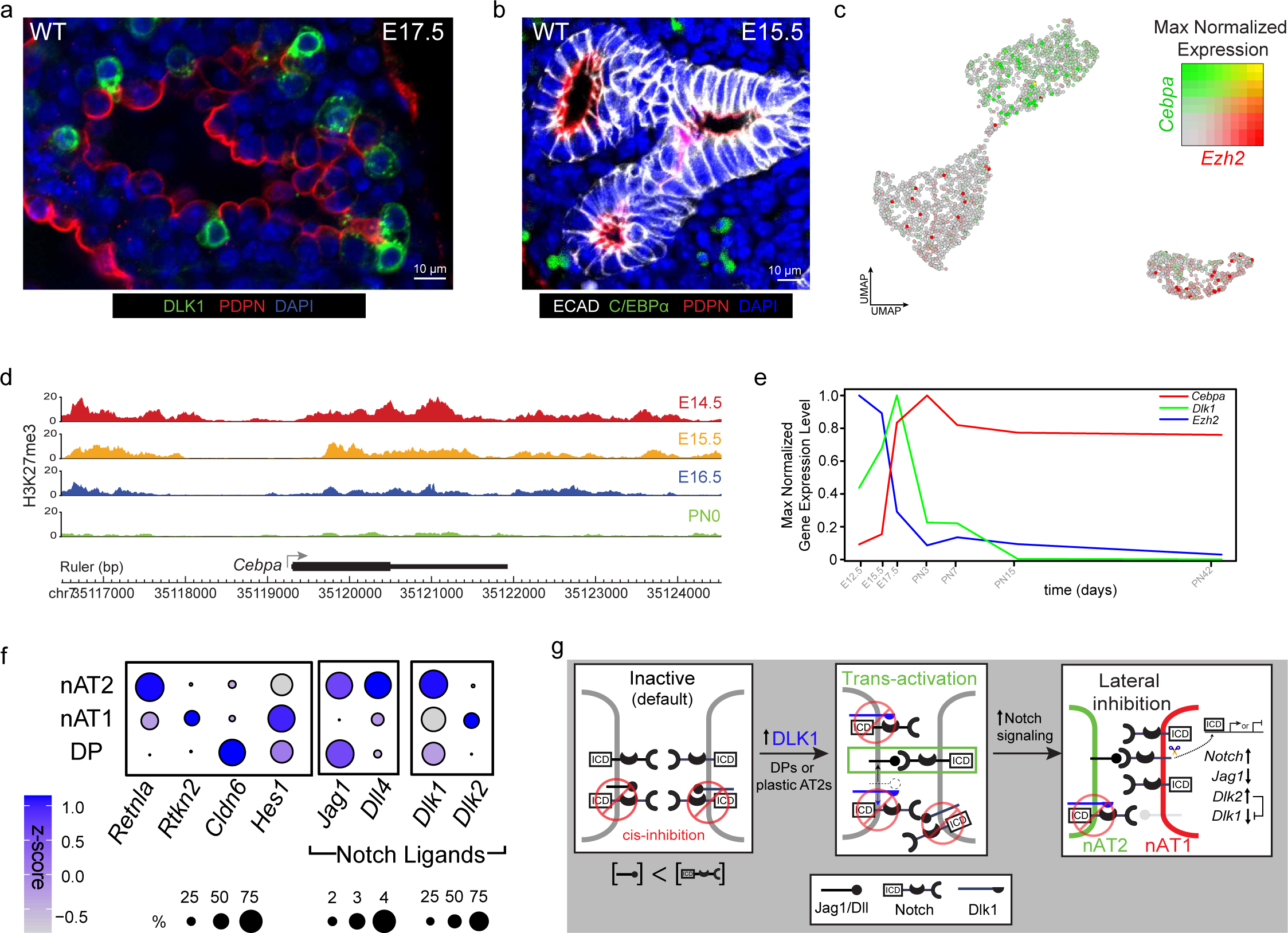
Embryonic expression of C/EBPα and DLK1. **a** Distal epithelial branch of an e17.5 lung immunostained for DLK1 (green), PDPN (red), and DAPI (blue). Bar, 10 µm. **b** Distal epithelial branch of an e15.5 lung immunostained for C/EBPα (green), PDPN (red), E-cadherin (ECAD, white), and DAPI (blue). No C/EBPα^+^ cells are detected in the distal epithelial branches at this point in development. Bar, 10 µm. **c** *Cebpa* (green) and *Ezh2* (red) co-expression on scRNAseq UMAP of the developing lung^33^ shows mutually exclusive expression. **d** Analysis of ENCODE epigenetic dataset mapping of the PRC2 modification H3K27me3 across developmental timepoints (e14.5 – P0) at the *Cebpa* locus. **e** Max normalized gene expression for *Cebpa* (red), *Dlk1* (green) and *Ezh2* (blue) during development from scRNAseq data of the developing lung^37^, showing the relation of the components forming the I2-FFL that generates a pulse of *Dlk1*.**f** Dot plot showing expression of the Notch signaling pathway genes (ligands and its downstream activation) have distinct pattern as nascent cell fates are specified. **g** schematics of our working model how a Dlk1 pulse generator regulate AT2’s accessibility to fate plasticity via Notch signaling.

**Supplementary Figure 7:**
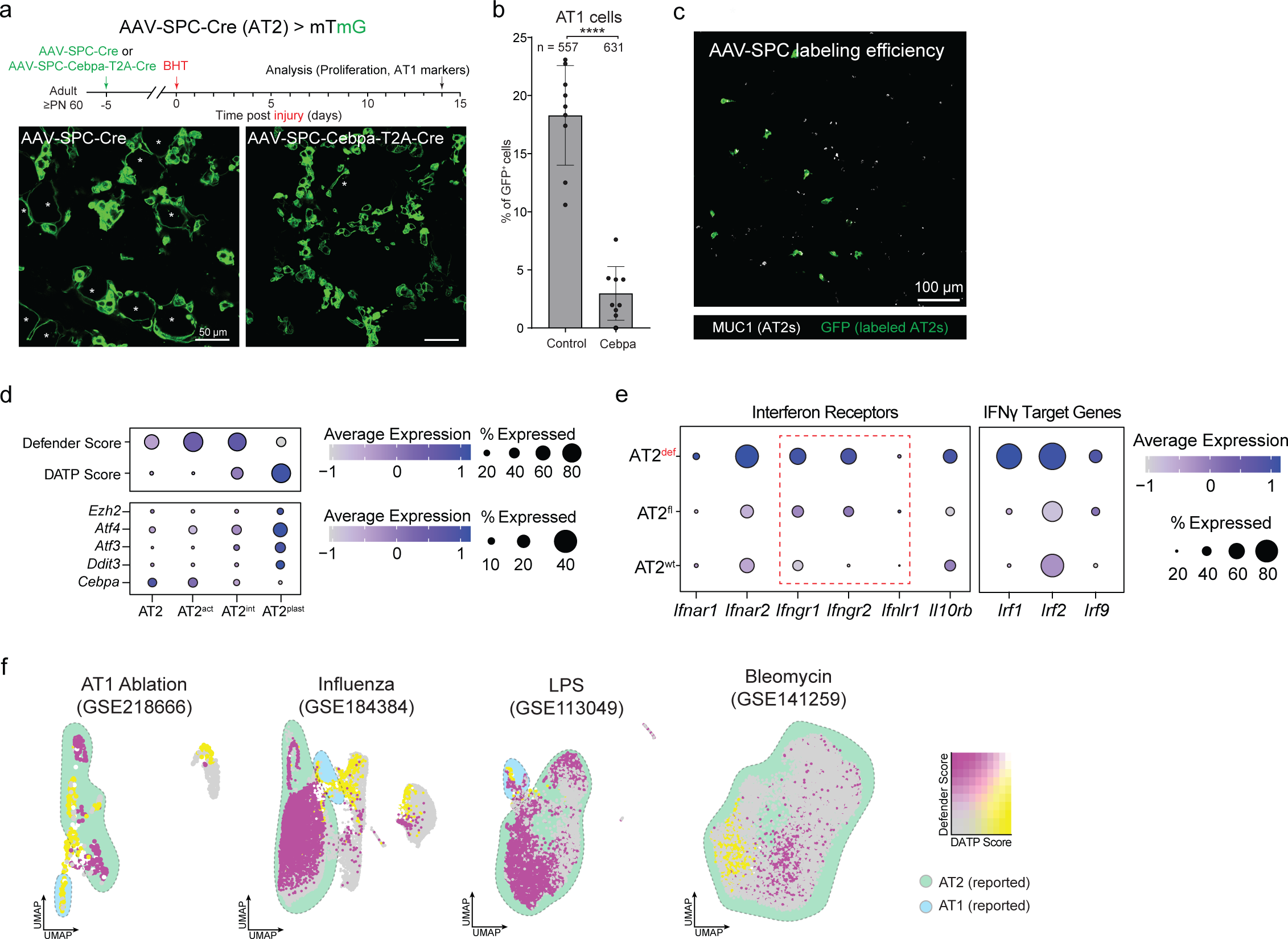
AT2 defensive state and interferon signaling. **a** Experimental design (top). 5 days after AAV-SPC-Cre (Control; left) or AAV-SPC-Cebpa-T2A-Cre (right) instillation into lungs of adult *Rosa26^mTmG^* mice to induce constitutive GFP-Cre expression and overdrive expression of *Cebpa* in AT2 cells, lungs were injured by BHT (225 mg/kg) intraperitoneal injection (i.p.). Lungs were isolated and immunostained for Cre reporter (GFP) and AT1 markers 14 days post injury (dpi). Immunostained images (bottom) show GFP^+^ AT1 cells (asterisk) in the control, but little to none when C/EBPα is overexpressed. Bars, 50 µm. **b** Quantification of (**a**) for percent GFP^+^ cells that differentiated into AT1 cells (*n* = 557 GFP^+^ cells sampled for the control condition and 631 for the *Cebpa* overdrive in experimental triplicate). \*\*\*\**p* = 6.5 × 10^-10^ (Student’s two-sided *t*-test, data as mean ± SD). **c** Immunostained adult lung showing GFP-labeling efficiency of AT2 cells (MUC1; white) by AAV-SPC-Cre (Figure 7a). Bar, 100 µm. **d** Dot plot showing interferon receptors (left) and interferon-γ target genes (right) for the AT2 clusters from the scRNAseq analysis. The AT2^Def^ cluster is highly enriched in both receptors and target genes. Interferon-γ specific receptor genes (dashed red box) are upregulated in the AT2^Def^ state upon loss of *Cebpa*. **e** Dot plot showing expression of DATP and Defender gene scores (top), along with expression of *Cebpa*, *Ezh2*, and DATP-associated genes (*Atf3*, *Atf4*, *Ddit3*) (bottom) in the clusters identified from the bleomycin injury timecourse scRNAseq dataset^20^. **f** DATP- and Defender-associated gene score co-expression shows a mutually exclusive pattern in four other lung injury model besides Cebpa deletion model (Fig 7j).

## Notes

### Competing Interest Statement

The authors have declared no competing interest.

### Summary of Updates

Author list has been updated to add author onto manuscript.

