## Supplementary Tables 1-4 for "A molecular circuit regulates fate plasticity in emerging and adult AT2 cells"

Supplementary Table 1

| DP Score | AT1 Score | AT2 Score | nAT2 Score | mAT2 Score | Defender Score | DATP Score |
|---|---|---|---|---|---|---|
| Cldn6 | Pdpn | Sftpa1 | Dlk1 | Lyz2 | Itih4 | Cldn4 |
| Fbn2 | Cav1 | Sftpa2 | Ly6c1 | Lyz1 | Lcn2 | Sprr1a |
| Igfbp5 | Rtkn2 | Sftpb | Ly6a | Fabp5 | H2-Aa | Areg |
| Top2a | Hopx | Sftpc | Ly6e | Il33 | Chil1 | Edn1 |
| Mki67 | Gprc2a | Sftpd | Sftpd | Lamp3 | Iigp1 | Cyr61 |
| Ccnd2 | Emp2 | Lamp1 | Sftpa1 | Abca3 | Cd74 | Plaur |
| Ube2c | Cldn18 | Lamp3 | Arg2 | Fasn | H2-K1 | Tnip3 |
| Nasp | Clic5 | Abca3 | Lrg1 | Lrp2 | H2-Eb1 | Hbegf |
| Hmga2 | Ager | Nkx2-1 | Mt1 | Sftpc | Igtp | Cdkn1a |
| Cenpa | Aqp5 | Muc1 | Fth1 | Lrrc8c | Hp | S100a6 |
| Igfbp2 | | | Retnla | H2-Aa | Ifitm3 | S100a10 |
| Nop58 | | | Col6a2 | H2-Ab1 | H2-Q7 | Lgals3 |
| Hn1 | | | Tcfcp2l1 | H2-Eb1 | B2m | Tnfrsf12a |
| Fubp1 | | | Spink5 | Cyp2b10 | Gm4951 | Fn1 |
| Smc2 | | | Nupr1 | Gm10801 | Gbp2 | Ctgf |
| Mcm7 | | | Vnn1 | Inmt | Gbp5 | Itgb6 |
| Mcm3 | | | Pon1 | Cyp4b1 | H2-Ab1 | |
| Psat1 | | | Rab27a | Hpgd | Gm4841 | |
| Bzw2 | | | Scnn1a | Enpep | | |
| Snrpd1 | | | Gdpd2 | Ly6c1 | | |
| Mcm6 | | | Tspan2 | Ly6a | | |
| | | | Kcnj15 | Acoxl | | |
| | | | Lrrk2 | Tmem100 | | |
| | | | Glrx | H2-DMb1 | | |
| | | | Vldlr | Npw | | |
| | | | Cebpd | Mme | | |
| | | | Pmvk | Scgb1a1 | | |
| | | | Tspan11 | Calcrl | | |
| | | | Atp6v1c2 | Lrg1 | | |
| | | | Slc43a2 | Gstt1 | | |
| | | | Cds1 | | | |
| | | | Gif | | | |
| | | | Vnn3 | | | |
| | | | Tspan1 | | | |
| | | | Sftpb | | | |
| | | | Sgpp2 | | | |
| | | | Isca1 | | | |
| | | | Pon3 | | | |
| | | | Aqp1 | | | |
| | | | Lpl | | | |

Supplementary Table 2

| **nAT2 markers** (pct.nAT2>25%, log2FC ranked) | |
|---|---|
| top1-30 | top 31-60 |
| Cela1 | Dhrs7 |
| Ggt5 | Tmem213 |
| Mirg | Tuba8 |
| Meg3 | Mlph |
| Sec1 | Areg |
| Retnla | Gm11744 |
| Egln3 | Lipa |
| Qpct | Fabp12 |
| S100a9 | Ntn5 |
| Pglyrp1 | Tyrobp |
| BC100530 | Il34 |
| S100a8 | Ndufa4l2 |
| Filip1 | Ctsc |
| Dlk1 | Cygb |
| Atp6v0a4 | Rps6ka4 |
| Ncf1 | Rab27b |
| Klk13 | Rmdn2 |
| Isca1 | Clmn |
| Gnmt | Fcer1g |
| Inpp4b | Cd164l2 |
| Phgr1 | Bcl2a1b |
| Ass1 | Cebpa |
| Gjb6 | Ctsh |
| Snx29 | Hif3a |
| Vnn1 | Pex26 |
| Lbp | Nkd1 |
| Acss2 | Mitf |
| Tspan1 | Ankrd37 |
| Ndrg1 | Insig1 |
| Bcl2a1a | Ttc38 |

Supplementary Table 3

| Figure 4a screening list | | | |
|---|---|---|---|
| **DP (1-30)** | **DP (31-60)** | **nAT2** | **mAT2** |
| Sox9* | Mki67ip | Etv5* | Nupr1* |
| Etv5* | Smarca1 | Tcfcp2l1 | Etv5* |
| Tead2 | Bcl11a | Nupr1* | Ifnar2 |
| Mcm7 | Hmgn2 | Elf5* | Nr1d1 |
| Mcm6 | Carm1 | Cebpa* | Cebpa* |
| Mcm3 | Drg1 | Creb3l1 | Ehf |
| Hmgb3 | Egr2 | Rab15* | Rab15* |
| Mcm5 | Ndn | Sox9* | |
| Ilf3 | Nfyb | Rps6ka4 | |
| Elf5* | Trip13 | Per3 | |
| Ezh2 | Bmyc | | |
| Mcm4 | Mtf2 | | |
| Mcm2 | Whsc2 | | |
| Uhrf1 | Nfya | | |
| Basp1 | Polr2i | | |
| Dnmt1 | Taf6 | | |
| Myef2 | Foxm1 | | |
| Sox12 | Mllt3 | | |
| Rab15* | Tgif2 | | |
| Mybbp1a | Trp53 | | |
| Cbx2 | Gtf3a | | |
| Tfdp1 | Gtf2e2 | | |
| Rbm14 | Sap30 | | |
| Notch3 | Polr2j | | |
| Gli3 | E2f3 | | |
| Cbfb | Ash2l | | |
| Notch1 | Cdk2 | | |
| Ruvbl1 | Ruvbl2 | | |
| Plagl2 | Pdcd11 | | |

*Genes are ranked across multiple lists.

Supplementary Table 4

| Gene Ontology | -Log P Value | Fold Enrichment |
|---|---|---|
| defense response (GO:0006952) | 17.46 | 12.41 |
| response to type II interferon (GO:0034341) | 14.05 | 65.64 |
| response to external biotic stimulus (GO:0043207) | 12.88 | 10.74 |
| response to biotic stimulus (GO:0009607) | 12.70 | 10.43 |
| defense response to other organism (GO:0098542) | 13.06 | 13.22 |
| antigen processing and presentation of exogenous peptide antigen (GO:0002478) | 12.74 | 100 |
| response to other organism (GO:0051707) | 12.89 | 10.76 |
| antigen processing and presentation (GO:0019882) | 12.37 | 63.95 |
| antigen processing and presentation of exogenous antigen (GO:0019884) | 12.40 | 100 |
| biological process involved in interspecies interaction between organisms (GO:00444 | 12.30 | 9.78 |
| innate immune response (GO:0045087) | 11.92 | 16.09 |
| antigen processing and presentation of peptide antigen (GO:0048002) | 11.54 | 82.29 |
| response to organic substance (GO:0010033) | 11.55 | 6.51 |
| response to cytokine (GO:0034097) | 11.38 | 14.45 |
| response to stress (GO:0006950) | 11.20 | 5.45 |
| cellular response to organic substance (GO:0071310) | 10.15 | 8.03 |
